# *Bordetella* spp. block eosinophil recruitment to suppress lung iBALT formation

**DOI:** 10.1101/2022.09.27.509784

**Authors:** Nicholas J. First, Amparo Martínez, África González-Fernández, Sushma Bharrham, Connor Roan, Emily Cox, Jian Wang, Rona Scott, Matthew Woolard, Monica C. Gestal

## Abstract

A characteristic that differentiates pathogenic and opportunistic bacteria is that pathogens have been selected by their ability to suppress host inflammatory responses allowing colonization and persistence. *Bordetella* spp. are respiratory pathogens characterized for the arsenal of mechanisms they use to manipulate host immune responses. We have previously characterized a *B. bronchiseptica* mutant, RB50Δ*btrS*, that is not able to suppress host immune responses, resulting not only in rapid clearance of the infection but also long-term lung sterilizing immunity against reinfection with the three classical *Bordetella* spp. Interestingly, this strong immune response requires eosinophils. In this work our results indicate that wildtype *B. bronchiseptica*, RB50, blocks eosinophil pro-inflammatory functions to prevent the rapid recruitment of B and T cells to the lung that results in iBALT formation. Moreover, eosinophils promote a TH17 microenvironment within the iBALT that might be responsible for the long-term robust protective immunity generated by infection with this mutant. Overall, this work provides a novel role for eosinophils as promoters of adaptive immune responses and protective immunity, while also indicating that bacteria actively manipulate those cells to promote long-term persistence and reinfection.

## INTRODUCTION

During the evolutionary process, pathogens have been selected by their ability to suppress host immune responses to facilitate colonization and persistence. *Bordetella* spp. are respiratory pathogens that cause the disease known as whooping cough, a long-term persistent infection characterized for causing a high rate of mortality in infants and chronic disease in teens and adults^1,2^. In neonates, the development of pneumonia is critical because from there the disease progresses to pulmonary hypertension, that in approximately 32% further continue to fatal pertussis diseaseg^3^. One important aspect of *Bordetella* spp. success relies on an arsenal of mechanisms these bacteria possess to evade, suppress, and manipulate host immune responses^4–7^, specially phagocytic cells^8–11^. Understanding the molecular mechanisms that *Bordetella* spp. utilize to suppress host immunity and to initiate fatal pneumonic disease, can guide us towards the discovery of novel vaccines and therapeutics.

The classical *Bordetella* spp. are comprised of three species. *B. pertussis*, a strictly human pathogen; *B. parapertussis*, which infects humans and sheep; and *B. bronchiseptica*, which causes disease in a wide variety of mammals^7,12^. *B. bronchiseptica* is the evolutionary ancestor of the other two^12–14^ and shares over 98% of the genetic content. Because *B. bronchiseptica* causes a disease in mice that mimics the human pertussis illness in humans, it is a robust and reliable model of infection that allows mechanistic studies of host pathogen interactions at the molecular level^15^. Investigating *Bordetella* spp. mechanisms to suppress the host immune response led to the discovery of the sigma factor, *btrS*, a highly conserved regulator of bacterial immunosuppressive pathways. When mice were challenged with a mutant of *B. bronchiseptica* lacking *btrS* (RB50Δ*btrS*), infection was rapidly cleared from the lungs^16^. Moreover, mice generated robust protective immunity against reinfection with *B. bronchiseptica* but also to the human pathogens, *B. pertussis* and *B. parapertussis*^17^. Importantly, the protective immunity generated by this mutant lasted for at least 15 month, time at which lung sterilizing immunity was still robust against infection with the three *Bordetella* spp. due to an enhanced T helper (TH)17 memory response^17^. Investigating which immune cell population was critical for the generation of the robust protective TH17 immune response, we discovered an important role for eosinophils. In the absence of eosinophils, infection with RB50Δ*btrS* persists in the lungs for nearly as long as the infection caused by the wild-type bacteria^18^, suggesting a role for eosinophils during adaptive immune responses.

This type of granulocytes is well known in the context of parasitic infections and allergic and asthmatic reactions^19^. However, the role of eosinophils during health and disease has been long questioned by many researchers^20–22^. Associations between eosinophils, eosinophilic disorders, and *Bordetella* spp. infection were noted as early as 1970^23–25^. However, it took until 1978 that the bactericidal activity of eosinophils was reported^26^. Little was done to follow up this finding until relatively recently, when the role of eosinophils during *Helicobacter pylori* infections gained attention^27^. A very interesting observation was that during *H. pylori* infection, eosinophils promote maintenance of mucosal associated lymphoid tissue via A PRoliferation-Inducing Ligand (APRIL)^28^. Our previous results also suggested that, during respiratory infection with a *Bordetella bronchiseptica btrS*-null mutant, eosinophils contributed to the formation of lymphoid aggregates adjacent to lung bronchi^18^, further suggesting a role for these cells during adaptive mucosal responses.

We have now focused our research on investigating the role of eosinophils during the generation of TH17 sterilizing protective immunity. We used two different mouse models: the BALB/c and the eosinophil deficient ΔdblGATA-1^29^ (with a mutation in the GATA-1 gene) and as secondary model, we used C57BL/6 compared with the eosinophil deficient EPX/MBP^-/-^ mice^30^ (knockout for the eosinophil peroxidase gene). In this work, we have discovered that *Bordetella* spp. promotes persistence by blocking the formation of inducible bronchus-associated lymphoid tissue (iBALT) by suppressing eosinophil proinflammatory functions. In our studies using the wild-type *B. bronchiseptica*, RB50, which suppresses host immune responses, and the mutant RB50Δ*btrS*, which cannot suppress host immunity, we find that during infection with the RB50Δ*btrS*, but not with the RB50 strain, eosinophils promote TH1/TH17 responses that are localized within the iBALT, possibly through the upregulation of chemokines and cell adhesion molecules needed for lymphocyte trafficking and adhesion. Overall, our work suggests a novel role for eosinophils in lung infections that shifts the classical view of these cells as merely TH2 hyper-reactive cells to cells that are capable of promoting strong localized TH17 responses that allow for faster clearance and possibly better protective immunity^17^.

## RESULTS

### RB50Δ*btrS*-null promotes significant transcriptional changes in the BALB/c mice in response to infection

Our previous research indicates that the immune response to infection with a mutant strain of *B. bronchiseptica, RB50ΔbtrS*, is stronger than that triggered by the wildtype, RB50^17^. Moreover, infection with RB50Δ*btrS* promotes the development of protective immunity by enhancing TH1/TH17 mucosal responses that last for at least 15 months post-infection^17^, leading us to hypothesize that *btrS* blocks host inflammatory signaling cascades required for lymphocyte recruitment to promote the characteristic long-term persistence infection of *B. bronchiseptica*. To test this hypothesis, we first selected a time point (7 days post-challenge) during which bacterial burden in the lungs is similar between both strains (bacterial strains as well as mouse strain). Thus, potential differences that we may find are not the result of different bacterial load in the lungs, but due to the host immune response. Using RNA sequencing of transcripts extracted from whole lungs, we determined transcript abundance in lungs of BALB/c mice infected with RB50 or RB50Δ*btrS* strains to identify specific signatures that correlate with enhanced immune response and rapid bacterial clearance.

Interestingly, we found specific immune signatures for each infection. Principal Component Analysis (PCA) (**Figure S1**) revealed a trend for samples of the same type of infection to cluster together. That indicated that we could identify specific transcriptomic signatures that were enhanced following infection with each bacterial strain. We decided to restrict our analysis to those genes that presented a significance value of 0.05 or greater. A total of 1513 differentially expressed (DE) genes were identified in BALB/c infected with the wild-type strain RB50 (**Figure S1B**) (compared to uninfected mice). Most of the differentially expressed genes appeared to be up-regulated (**Figure S1C**). Mice infected with the RB50Δ*btrS* strain presented 3231 differentially regulated genes (compared to uninfected controls) (**Figure S1B**) where again, most of the differences were found on genes being upregulated in comparison to the uninfected controls (**Figure S1D**). A similar pattern was observed with the second mouse model, C57BL/6 (**Figure S2**).

Pathway analysis revealed that mice infected with RB50 presented upregulation of pathways related to *leukocyte migration and mediated immunity, humoral response, and adaptive immune responses* (**Figure 1A**) suggesting that, unsurprisingly, infection triggers an immune response. Pathways upregulated after RB50Δ*btrS* infection are indicative of more robust *acute and innate immune responses*, stronger *leukocyte and lymphocyte responses, more active antigen presentation and complement activation, as well as upregulation of T and B cell activation* (**Figure 1B**). Overall, our results suggest that infection with RB50Δ*btrS* triggers stronger innate and adaptive immune responses than RB50 infection. Excitingly already at day 7 post-infection, markers for adaptive response were significantly up-regulated suggesting that the mutant RB50Δ*btrS* might promote earlier adaptive immune responses than we thought based on our previously published data at day 14 post-infection^16^.

**Figure 1:**
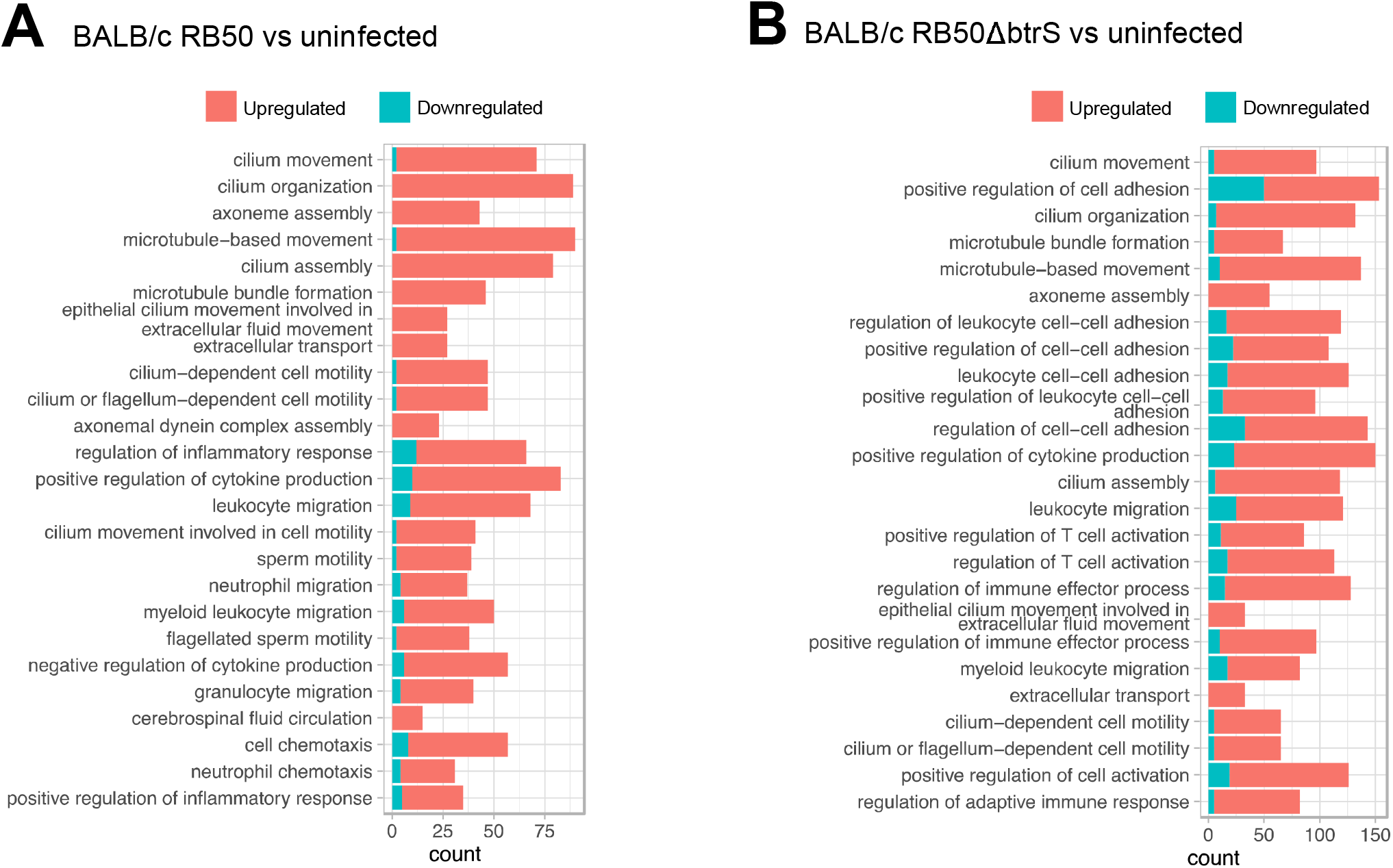
Infection with RB50Δ*btrS* promote the differential expression of genes related with adaptive inflammatory responses. RNA sequencing of the whole lungs was performed in 4 uninfected controls, 3 mice infected with RB50, and 4 mice infected with RB50Δ*btrS*. Over-representation analysis of BALB/c infected with RB50 (A) or RB50Δ*btrS* (B) differentially expressed genes. Top 25 p-adjusted pathways are illustrated and ordered by significance. Bar plots represent the number of DE genes participating in each pathway, whose expression is upregulated (red) or downregulated (blue) compared to uninfected control (n= 3-5 mice per group).

### RB50Δ*btrS* promotes adaptive immune responses

Our transcriptomic data revealed that RB50Δ*btrS* activates adaptive immune responses better than RB50. We have previously shown that RB50Δ*btrS* promotes recruitment of B and T cells to the lungs at day 14 post-infection^16^ and long-lasting protective immunity^17^. We hypothesized that *btrS* suppresses early recruitment of adaptive immune cells to the lungs. A heatmap analysis of surface markers and cytokines characteristic for T cells showed that lungs infected with RB50Δ*btrS* had increased transcript levels of genes associated with CD4 and CD8 T cells, including but not limited to TNFα, IL6, INFγ, and IL17, in line with our pathway analysis. While we also detected higher levels of TNFαand IL6 in RB50 infected lungs compared to uninfected controls, the changes were not quite as pronounced. No significant increase was observed for CD8 markers. B cell markers were only marginally increased after infection with RB50Δ*btrS* but not RB50(**Figure 2A**). Flow cytometry gating for live cells, followed by CD45^+^, and then we selected CD3^+^ and CD19^+^ confirmed increased numbers of T cells in RB50Δ*btrS* infected lungs at 7 dpi compared to RB50 infected lungs. We also observed larger numbers of B cells in lungs of RB50Δ*btrS* infected Balb/c mice (**Figure 2B**). Essentially identical results were obtained in Balb/c and C57BL/6 mice (**Figure 2C**). Overall, our results indicate that RB50Δ*btrS* promotes a better T and B cell response than Rb50 at day 7 post infection.

**Figure 2:**
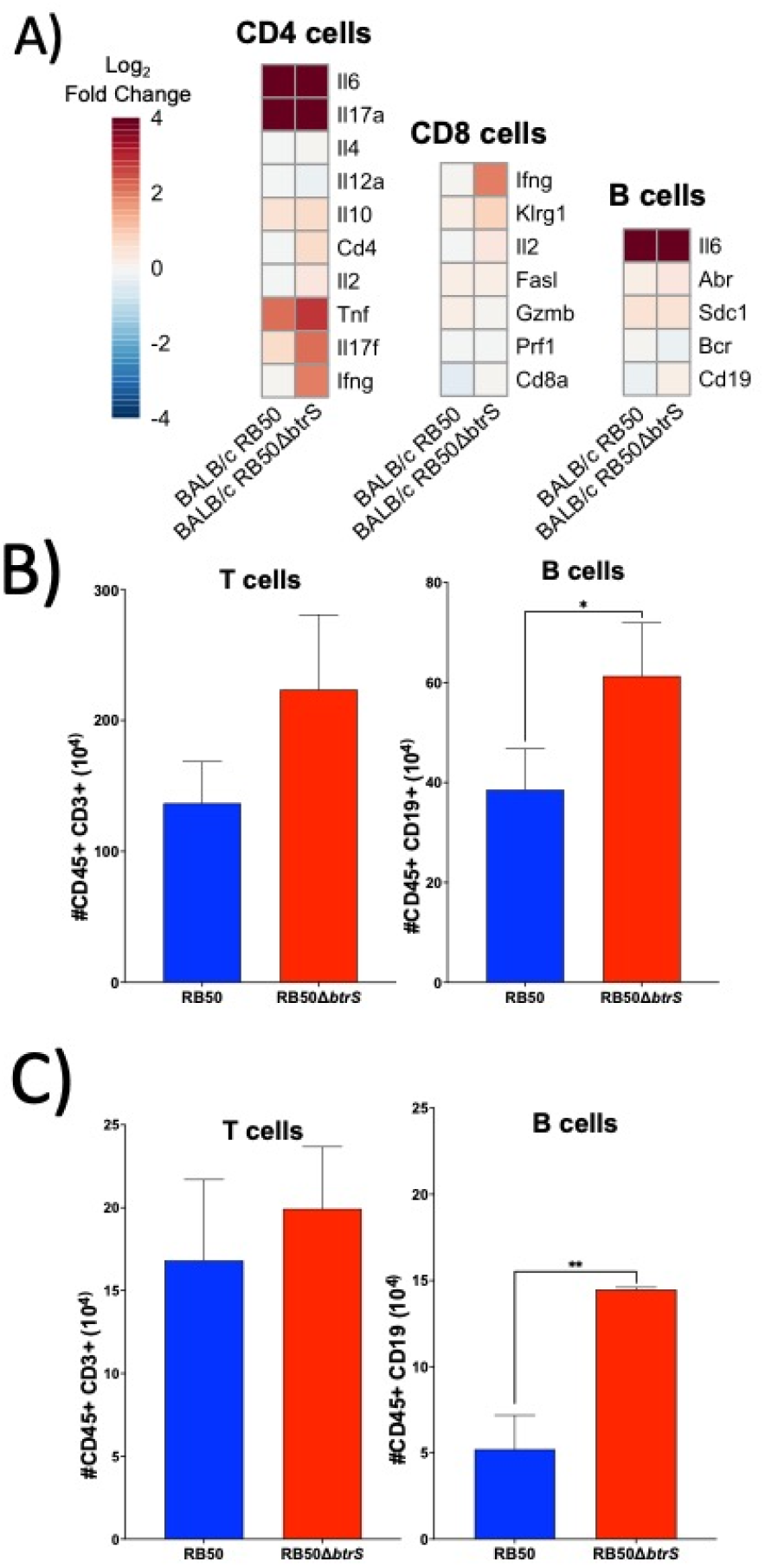
*btrS* suppress recruitment of B and T cells to the lungs. Mice were uninfected or intranasally challenged with 30 μl of PBS containing 5×10^5^ RB50 or RB50Δ*btrS*. At day 7 post infection mice were euthanized and lungs were collected in cold PBS. (A) Following RNA sequencing of the lungs, genes correlated with CD4, CD8 and B cell markers were plotted on a heatmap where darker read represents more expression and darker blue represents less expression (n= 3-5 mice per group). (B) Single suspension of lungs cells of BALB/c mice infected with RB50 (blue) or RB50Δ*btrS* (red) was utilized to perform flow cytometry staining (n=8-9 mice per group). (C) Single suspension of lungs cells of C57BL/6J mice intranasally infected with RB50 (blue) or RB50Δ*btrS* (red), was utilized to perform flow cytometry staining (n=3-5 mice per group). Two-Way ANOVA was performed to analyze the data. * p<0.05 and ** p<0.01.

To validate our findings, we measured transcript levels of representative genes by qRT-PCR including Interleukin (IL)17, interferon gamma (INFγ), and Chemokine (C-C motif) ligand 19 (CCL19). Our results showed that these genes were significantly increased in the two independent animal models, BALB/c and C57BL/6J, after infection with RB50Δ*btrS* but not with RB50 (**Figure S3**). Overall, these results indicate that challenge with RB50Δ*btrS* promotes recruitment of B and T cells indicative of enhanced adaptive immune responses.

### BtrS suppresses Th17 mucosal responses

Our previous research revealed that in comparison to the wildtype RB50 infection, the mutant RB50Δ*btrS* bacteria induce long-lasting Th17 mucosal responses^16,17^. It has been previously demonstrated that TH17 responses are critical for *Bordetella* spp. clearance^16,31^ and also for the generation of protective immunity^17,32,33^. We analyzed the above-mentioned transcriptomic data focusing on TH17 mucosal responses. We observed higher transcript levels of TH17 related cytokines in RB50Δ*btrS* compared to RB50 (**Figure 3A, S3**). To confirm these findings, we isolated lung T cells at day 7 post-infection using negative CD3 selection magnetic beads. After isolation, we stimulated them with CD3/CD28 for 48 h and performed an unbiased secretome analysis using the Isoplexis platform (**Figure 3B**). In accordance with our previous results, we found higher levels of GM-CSF, IL17, INFγ, and IL2 after infection with RB50Δ*btrS* compared to RB50 (**Figure 3B**). We also subjected lung immune cells to intracellular flow cytometry staining and found an increase of INFγand IL17 in T cells from lungs of BALB/c mice infected with RB50Δ*btrS* but not RB50 (**Figure 3C**). The same results were obtained using our secondary model of C57BL/6J (**Figure 3D**). Taken together, our data indicate that *Bordetella bronchiseptica* blocks TH17 mucosal responses via a *btrS* dependent mechanism.

**Figure 3:**
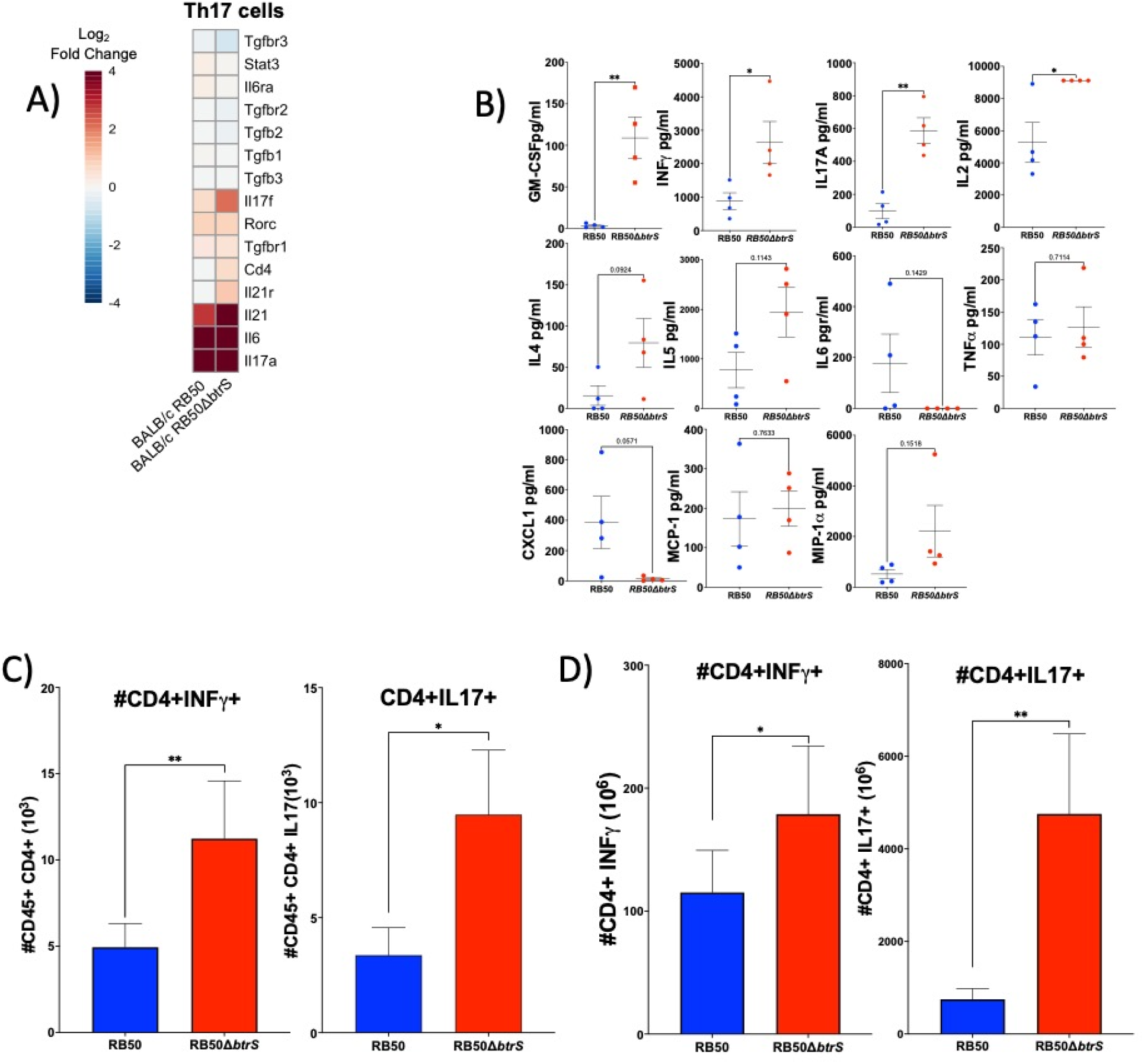
*btrS* suppress TH17 responses. Mice were uninfected or intranasally challenged with 30 μl of PBS containing 5×10^5^ RB50 or RB50Δ*btrS*. At day 7 post infection mice were euthanized and lungs were collected in cold PBS. (A) Following RNA sequencing of the lungs, genes correlated TH17 mucosal responses were plotted on a heat map where darker read represents more expression and darker blue represents less expression (n= 3-5 mice per group). (B) Lungs of BALB/c mice infected with RB50 (blue) or RB50Δ*btrS* (red) were used to obtain isolated T cells. After 48 hours stimulation with CD3/CD28 Dynobeads, the supernatant was collected to perform a Bulk CodePlex assay (n= 4 mice per group). (C) Single suspension of lungs cells of BALB/c mice infected with RB50 (blue) or RB50Δ*btrS* (red) was utilized to perform flow cytometry staining (n=8-9 mice per group). (D) Single suspension of lungs cells of C57BL/6J mice intranasally infected with RB50 (blue) or RB50Δ*btrS* (red), was utilized to perform flow cytometry staining (n=3-5 mice per group). Two-Way ANOVA was performed to analyze the data. * p<0.05 and ** p<0.01.

### Eosinophils are required for clearance of RB50Δ*btrS* but not RB50 and for recruitment of adaptive immune cells to infected lungs

We have previously shown that clearance of RB50Δ*btrS* is dependent on eosinophils^18^. Since we never explored the role of eosinophil in RB50 infection, we infected BALB/c and the eosinophil-deficient ΔdblGATA-1 mice with the RB50 strain and enumerated colonies at different times post-infection. We saw no significant differences in bacterial burden and the clearance dynamics of RB50 (**Figure S4A**). In contrast, we again observed a delayed clearance of RB50Δ*btrS* from the lungs of ΔdblGATA-1 mice (**Figure S4B**). Subsequently, we confirmed the importance of eosinophils for bacterial clearance using the second mouse model of eosinophil deficiency, EPX/MBP^-/-^, (**Figure 4A**). These results confirm the importance of eosinophils for clearance of RB50Δ*btrS* and, furthermore, suggest that RB50 blocks eosinophil effector functions via a *btrS* dependent pathway masking the pro-inflammatory effects of eosinophils during *Bordetella* spp. We also compared host transcript levels in the lungs of infected and uninfected ΔdblGATA-1 mice and saw no significant difference in pathway analysis and specific transcripts characteristic of CD4, CD8 and B cells when comparing RB50 and RB50Δ*btrS* infected mice (**Figure 4B-D** and **S2**). We also found similar numbers of T and B cells (**Figure 4E**) and confirmed these findings in the EPX/MBP^-/-^ mouse model (**Figure 4F**). Overall, our results suggest that the absence of eosinophils impair B and T cell recruitment ot he lungs of the RB50Δ*btrS* infected mice.

**Figure 4:**
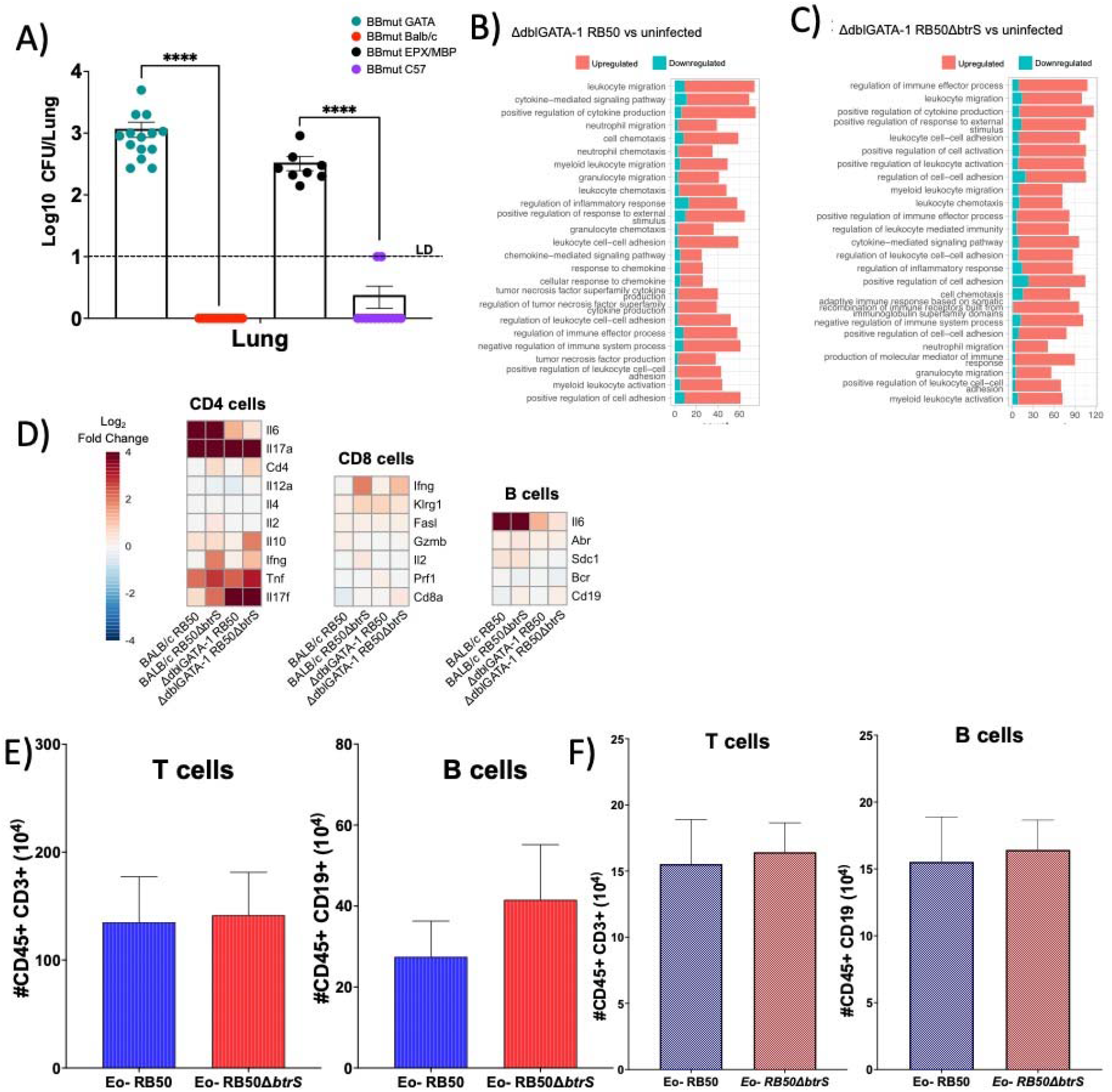
Eosinophils are required to promote B and T cell responses in the *btrS* null mutant. (A) BALB/c (green), ΔdblGATA-1 (red), C57BL/6J (black), and EPX/MBP^-/-^ (purple) mice, were intranasally inoculated with RB50Δ*btrS*. At day 14 post-infection, lungs were collected to enumerate colony forming units (CFU) in Bordet-Gengou Streptomycin (BGS) agar (n= 10-14 mice per group). (B, C, D) Over-representation analysis of ΔdblGATA-1 infected with RB50 (B) or RB50Δ*btrS* (C) differentially expressed genes. Top 25 p-adjusted pathways are illustrated and ordered by significance. Bar plots represent the number of DE genes participating in each pathway, whose expression is upregulated (red) or downregulated (blue) compared to uninfected control. (D) CD4, CD8 and B cell markers were plotted on a heatmap where darker read represents more expression and darker blue represents less expression (n= 3-5 mice per group). (E) ΔdblGATA-1 mice were intranasally infected with RB50 (blue) or RB50Δ*btrS* (red). At day 7 post-infection single cell suspension was used to perform flow cytometry staining. Total T and B cell numbers are show (n= 8-9 mice per group). (F) EPX/MBP^-/-^ mice were intranasally infected with RB50 (blue) or RB50Δ*btrS* (red). At day 7 post-infection single cell suspension was used to perform flow cytometry staining. Total T and B cell numbers are shown (n= 3-5 mice per group). Two-Way ANOVA was performed to analyze the data and **** p≤0.0001.

### Eosinophils promote Th17 mucosal responses after infection with RB50Δ*btrS*

Our previous results revealed a failure in IL17 secretion of ΔdblGATA-1 mice when challenged with RB50Δ*btrS*^18^, suggesting that absence of eosinophils might suppress Th17 responses. To test the hypothesis that the RB50Δ*btrS* infection promotes a Th17 mucosal response that requires eosinophils, we explored the expression levels of diverse specific genes related to the immune response in the lungs of ΔdblGATA-1 mice infected with either RB50 or RB50Δ*btrS* (**Figure 5A**). Our transcriptomic data indicated that in ΔdblGATA-1 mice following infection with RB50, Il17a and IL17f were increased. However, the rest of the markers that were previously upregulated in BALB/c, were decreased in the ΔdblGATA-1 mice, including IL6 and IL21. Similar results were observed for the RB50Δ*btrS* infected group. Overall, the transcriptomic data suggest that IL17 responses might be impaired in the eosinophil deficient mice.

**Figure 5:**
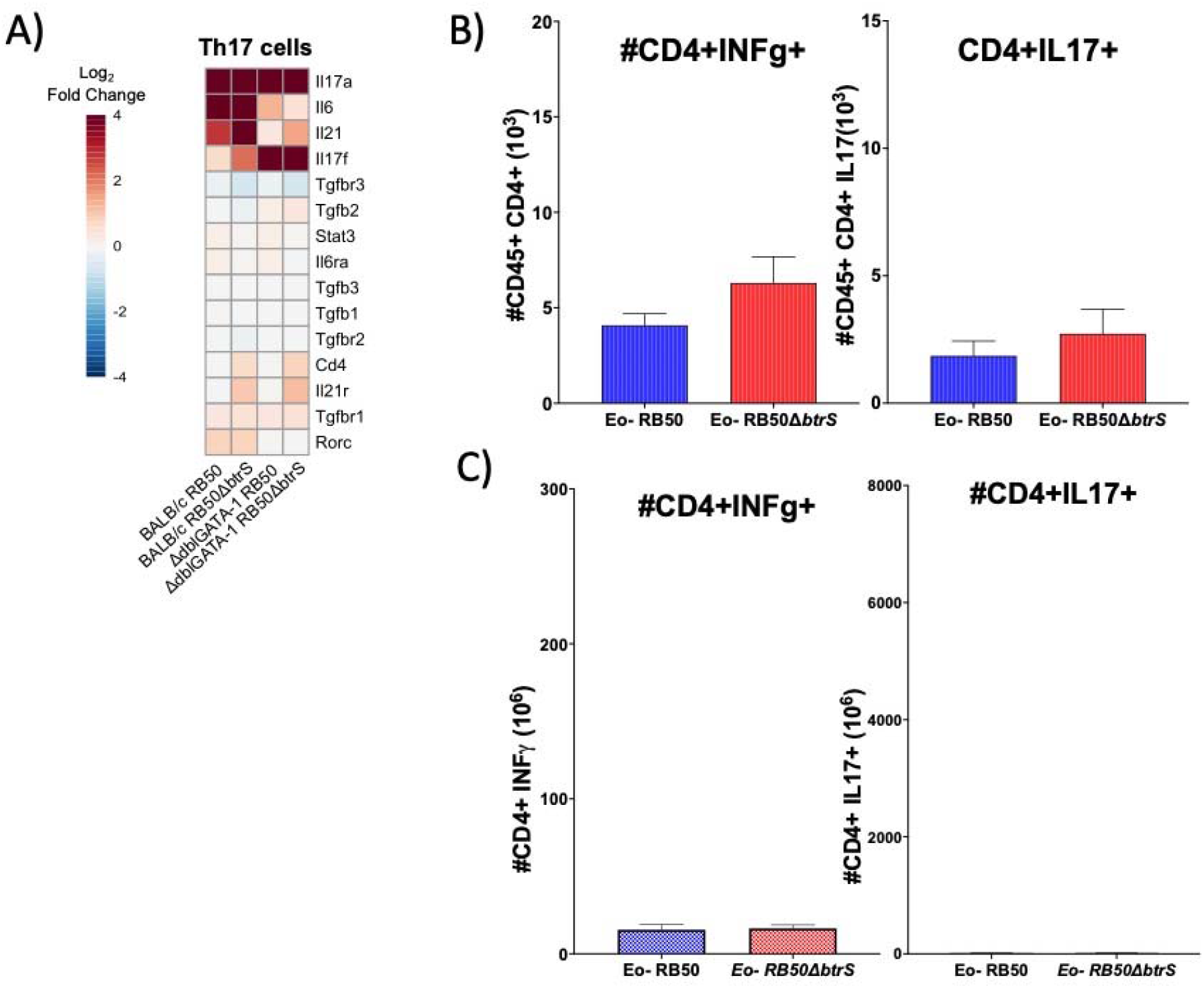
Eosinophils are required to promote TH17 responses in the *btrS* null mutant. (A) ΔdblGATA-1 mice were intranasally infected with RB50 or RB50Δ*btrS*. At day 7 post-infection RNA sequencing was performed. Heatmap representing expression levels of some genes Th17 responses. Color scale represent log2 fold change values. (n= 3-5 mice per group). (B) ΔdblGATA-1 mice were intranasally infected with RB50 (blue) or RB50Δ*btrS* (red). At day 7 post-infection single cell suspension was used to perform flow cytometry staining. Total CD4+ T cells were used to gate INFγand IL17 (n= 8-9 mice per group). (F) EPX/MBP^-/-^ mice were intranasally infected with RB50 (blue) or RB50Δ*btrS* (red). At day 7 post-infection single cell suspension was used to perform flow cytometry staining. Total CD4+ T cells were used to gate INFγand IL17. (n= 3-5 mice per group). Two-Way ANOVA was performed to analyze the data.

To follow up with these findings, an intracellular staining was performed to investigate the expression of INFγand IL17 in lung CD4 T cells of eosinophil deficient infected mice. In contrast to BALB/c mice infected with RB50Δ*btrS*, whose CD4 cells secreted significantly higher levels of INFγand IL-17, ΔdblGATA-1 mice revealed an impaired secretion of INFγand IL17 (Figure 5B), and the differences found between RB50 and RB50Δ*btrS* infected groups disappear in ΔdblGATA-1 mice. The same results were observed with the EPX/MBP^-/-^ mice (**Figure 5C**), were the cytokine production by CD4 T cells was severely impaired for both INFγand IL17 in comparison with the C57BL/6J wildtype. Thus, our results suggest that *Bordetella bronchiseptica* blocks Th17 mucosal responses via a *btrS* and eosinophil dependent mechanism.

### iBALT formation at day 7 post-infection with a *btrS* mutant strain of *B. bronchiseptica*

iBALT is known to be formed in the lungs in response to a range of pathogens and inflammatory stimuli, including gram-negative bacteria such as *Pseudomonas aeruginosa*^34,35^ and *Klebsiella pneumonia*^36^. A common feature of the iBALT during infection with these pathogens is that an initial lymphocyte accumulation can begin already on day 7^36^, leading us to the hypothesis that RB50Δ*btrS* could promote an early iBALT formation that leads to the robust protective immunity previously observed^17^.

To investigate iBALT signatures, we first looked at our transcriptomic data looking at genes that were differentially expressed and associated with the formation of germinal centers, including iBALT. The most upregulated hits were two genes whose expression is characteristic of high endothelial venules (HEVs): MAdCAM-1^37^ and GlyCAM-1^38^ (**Figure 6A**), as well as other important iBALT genes such as Fut7^39^, Lta^40^ and Ltb^41^, all of which have been reported to be influenced by IL17^41,42^. HEVs are specialized venules that mediate blood lymphocyte extravasation into lymphoid tissues. This list also included *Ccl19*, a pivotal chemokine in lymphocyte recruitment into lymph organs^43^, importantly, Ccl19 was increased in both mouse models following infection with the RB50Δ*btrS* strain (**Figure S3**). In conclusion, transcriptional changes suggest an increase in cell recruitment and adhesion after infection with the RB50Δ*btrS* strain, which can evoke iBALT formation.

**Figure 6:**
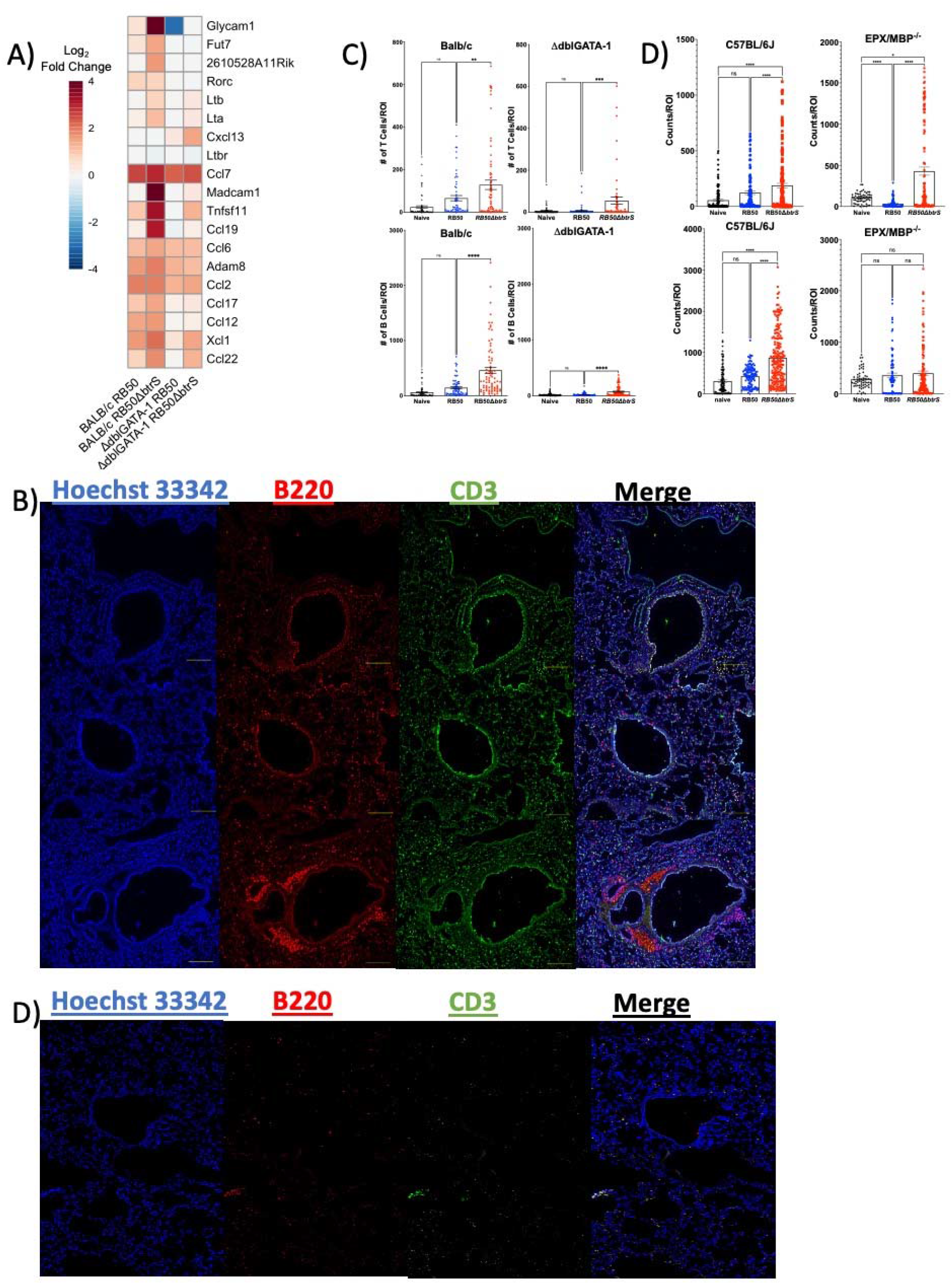
Eosinophils are required to promote iBALT formation during the *btrS* null mutant infection. (A) BALB/c and ΔdblGATA-1 mice were uninfected or intranasally infected with RB50 or RB50Δ*btrS*. At day 7 post-infection RNA sequencing was performed. Heatmap representing expression levels of the most representative genes related to iBALT. Color scale represent log2 fold change values. (n=3-5 mice per group). (B, C) BALB/c and ΔdblGATA-1 mice were intranasally infected with RB50 (blue) or RB50Δ*btrS* (red). At day 7 post-infection lungs were perfused and paraffin embedded. Three sections per mouse were stained with Hoechst (blue), CD3 to stain T cells (green), and B220 to stain B cells (red). Images were acquired in the Olympus CSU W1 Spinning Disk Confocal System 20x magnification. (n= 6 mice per group), 3 section per mice and 10 regions of interest. (C) Images were acquired and analyzed using Keyence software (n= 6 mice per group), 3 section per mice and 10 regions of interest. (D) C57BL/6J and EPX/MBP-/-mice were intranasally infected with RB50 (blue) or RB50Δ*btrS* (red). At day 7 post-infection lungs were perfused and paraffin embedded. (n= 6 mice per group), 3 section per mice and 10 regions of interest. (E) Three sections per mouse, ΔdblGATA-1, were stained with Hoechst (blue), CD3 to stain T cells (green), and B220 to stain B cells (red). ROI: Region of Interest (n= 6 mice per group), 3 section per mice and 10 regions of interest. Two-Way ANOVA was performed to analyze the data. * p≤0.05, ** p≤0.01, *** p≤0.001, and **** p≤0.0001.

To follow up with these findings, lungs of BALB/c and C57BL/6J mice challenged with RB50 or RB50Δ*btrS* were collected at day 7 post-infection to perform immunofluorescence staining of paraffin embedded tissues. We performed staining of three independent sections of 6 different BALB/c mice per group and acquired 10 different images per section per animal. Our results show that B and T cells numbers were low in the lungs of RB50 infected BALB/c mice, comparable to uninfected naïve controls (**Figure 6B** and **6C**). Moreover, those cells were scattered throughout the lung section. Contrary, in the RB50Δ*btrS* infected mice, B and T cells numbers were higher (**Figure 6C**), and those cells were forming aggregates characteristic of iBALT (**Figure 6B**). To further confirm that these aggregates correspond to iBALT, we performed a staining with a marker for endothelial lymphoid vessels. Our results revealed that in the BALB/c mice infected with the RB50Δ*btrS* strain, the aggregates were surrounded by endothelial hyaluronan receptor 1 (Lyve-1+) lymphatic vessels, a distinctive feature of iBALT (**Figure S5A**), and in fact the number of positive Lyve1 colocalizing with B cells was higher in the RB50Δ*btrS* infected group (**Figure S5B**), further confirming that these aggregates correspond to iBALT.

Similar results were obtained following infection of C57BL/6J mice with the RB50Δ*btrS* analyzed at day 7 post-infection (**Figure 6D**). However, in C57BL/6J mice the lymphocyte aggregates observed were smaller than in BALB/c (**Figure S5C**) similar to what has been previously reported^44^. Overall, our results show that lung B and T cells from RB50Δ*btrS* but not RB50 infected mice were forming iBALT, as early as day 7 post-infection; suggesting that *btrS* might suppress or delay iBALT formation.

### Eosinophils are required for iBALT formation during RB50Δ*btrS* infection

We have previously shown that eosinophil deficient Δdbl-GATA-1 mice display decreased numbers of B and T cells and delayed clearance^18^. Therefore, we hypothesized that RB50Δ*btrS* iBALT formation could be dependent on eosinophils. To test this hypothesis, immunopathological studies were performed using ΔdblGATA-1 and EPX/MBP^-/-^ infected with RB50 or RB50Δ*btrS*. Our results showed that the lungs of Δdbl-GATA-1 infected with RB50 had lower numbers of B and T cells than the uninfected control (**Figure 6C**), specially for B cells that were significantly reduced in ΔdblGATA-1 mice. Moreover, the few cells observed were scattered throughout the lung tissue and no aggregates were observed, not even for the RB50Δ*btrS* infected group (**Figure 6B**). Same results were obtained using the C57BL/6J and eosinophil deficient EPX/MBP^-/-^ pair (**Figure 6D** and **6C**). Although the numbers of B and T cells in the EPX/MBP^-/-^ mice were not as dramatically reduced as in the ΔdblGATA-1 mice (**Figure 6D** and **S5C**), the aggregates were still absent, further confirming a role for eosinophils in iBALT formation.

Overall, our results indicate that eosinophils are required for iBALT formation. Interestingly even our lymphatic endothelial marker was affected by the absence of eosinophils, suggesting that they are involved in many of the processes implicated in iBALT formation.

### iBALT localized Eosinophils secrete IL-17

Previous literature suggests that the formation of iBALT is dependent on IL-17 signaling^45^; as well as other chemokines and cytokines that we have seen increased after infection with RB50Δ*btrS* in our wildtype mice^46^. Based on our transcriptomic and flow cytometry data, we hypothesized that eosinophils localize within the iBALT and promote a Th17 microenvironment after challenge with the RB50Δ*btrS* strain.

To test this hypothesis, we first conducted an *in vitro* experiment where we challenged bone marrow derived eosinophils at a Multiplicity of Infection (MOI) of 10 with the RB50 or mutated RB50Δ*btrS* strains and performed an unbiased LegendPlex assay at 4 hours post-infection to evaluate secretion of IL17 from eosinophils in response to infection with these two bacterial strains (**Figure 7A**). Eosinophils infected with RB50Δ*btrS* but not RB50 secreted significantly higher levels of IL17 in comparison to the uninfected control, suggesting that eosinophils may contribute to an increased TH17 response.

**Figure 7:**
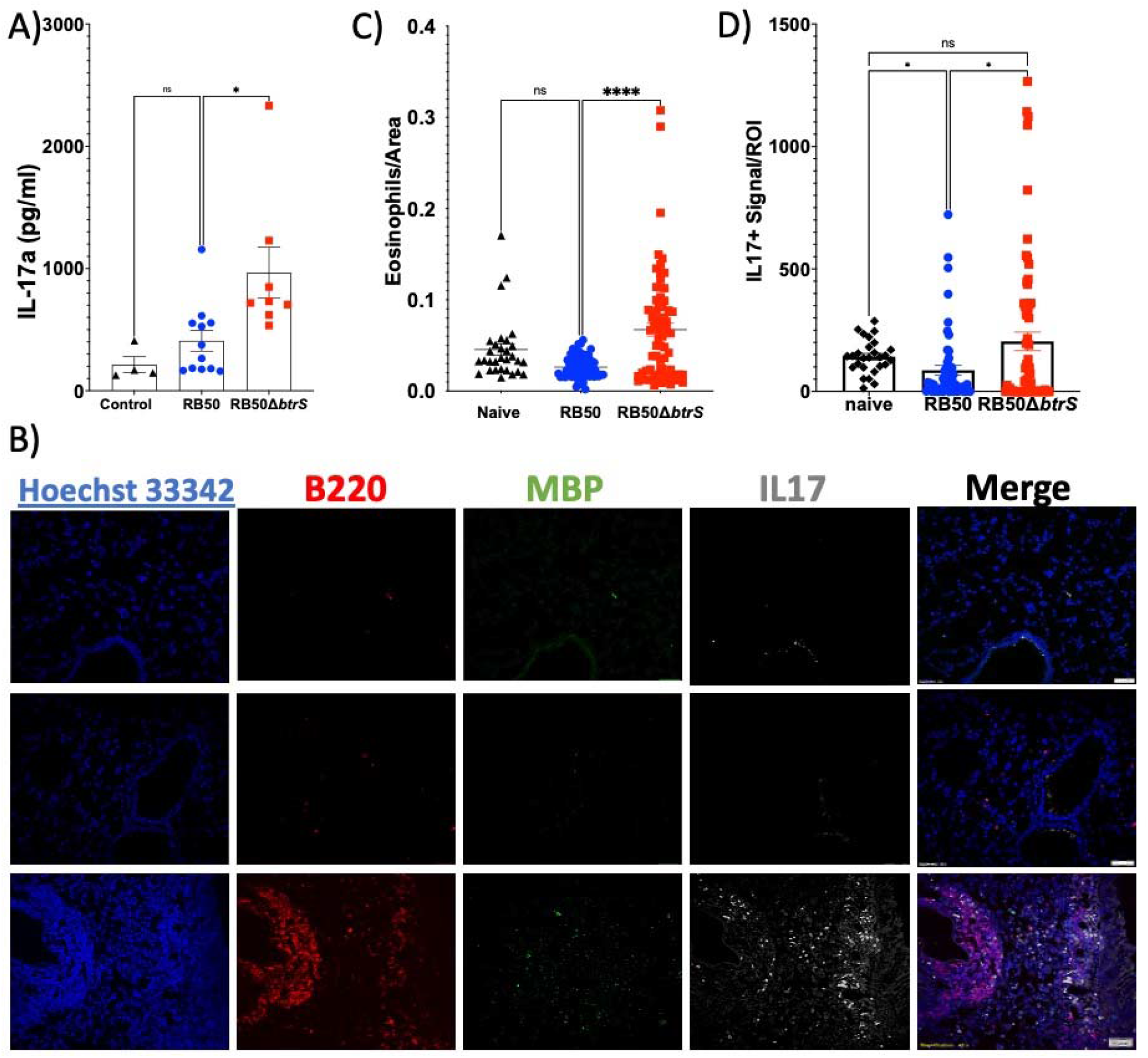
Eosinophils are required to promote a TH17 microenvironment within the iBALT formation during the *btrS* null mutant infection. (A) Eosinophils were uninfected (black) or challenged with RB50 (blue) or RB50Δ*btrS* (red) at an MOI of 10. At four hours post-infection the supernatant was collected to perform a LegendPlex assay. Each dot represents the average of two technical replicates. This experiment was performed in three individual biological replicates each of them containing 2-3 technical replicates. (B) BALB/c mice were intranasally infected with RB50 or RB50Δ*btrS*. At day 7 post-infection lungs were perfused and paraffin embedded. Three sections per mouse were stained with Hoechst (blue), B220 to stain B cells (red), MBP to stain eosinophils (green), and IL17 (grey). Images were acquired in the Olympus CSU W1 Spinning Disk Confocal System at 40x magnification. (n= 4 mice per group), 3 section per mice and 10 regions of interest. (C and D) BALB/c mice were intranasally infected with RB50 (blue) or RB50Δ*btrS*(red). At day 7 post-infection lungs were perfused and paraffin embedded. Three sections per mouse were stained with Hoechst, B220 to stain B cells, MBP to stain eosinophils, and IL17. Images were acquired and analyzed using Keyence software (n= 6 mice per group), 3 section per mice and 10 regions of interest., (C) Shows the number of eosinophils co-localizing with B cells. (D) Shows the number of positive IL17 signal co-localizing with B cells. ROI: Region of Interest. Two-Way ANOVA was performed to analyze the data. * p≤0.05 and **** p≤0.0001.

To follow up with the *in vitro* findings and based on our previous data indicating that the wildtype mice infected with RB50Δ*btrS* present stronger IL17 responses, we asked if the eosinophils are in the proximity of the iBALT. We sequentially immunostained lung section for B220+ (B cell), MBP+ (eosinophils), and IL-17A. We used only BALB/c mice because the eosinophil deficient mice do not have iBALT and the C57BL/6J have smaller aggregates. We used three sections of 6 different mice per group and acquired 10 regions of interest per section. While eosinophils were scattered in the lungs of RB50 infected mice (**Figure 7B** and **7C**); eosinophils co-localized near the B cells within the iBALT areas in mice challenged with RB50Δ*btrS* strain (**Figure 7B** and **7C**). Even though the IL17 signal was higher in the RB50 infected lungs than in the uninfected mice, it was scattered and did not colocalize with the iBALT (**Figure 7B**). The lungs of mice infected with RB50Δ*btrS* also presented more IL17 signal (**Figure 7D**) which colocalized with eosinophils and B cells in the iBALT (**Figure 7B**). Thus, our results indicate that eosinophils colocalize within the iBALT after challenge with the RB50Δ*btrS* strain promoting a Th17 microenvironment that facilitates rapid clearance^16^ of the RB50Δ*btrS* strain and possibly is responsible of the long-lasting immunity generated by this strain^17^.

## DISCUSSION

A current challenge in the field of vaccine development relies on the limited generation of mucosal protective immunity. Many researchers are focusing on improving the current mucosal vaccines by using live attenuated vaccines^47,48^, outer membrane vesicles^49,50^, adding novel antigenic targets^51^ or improving intranasal delivery techniques^52^. However, complete sterilizing immunity is still not achieved with the current platforms. Our results reveal that TH2 cell populations, such as eosinophils, play pivotal roles in the generation of long-lasting lung sterilizing immunity against *Bordetella* spp. infection. We for the first time propose a role for eosinophils in the generation of TH17-iBALT in the lungs responsible for rapid clearance and possibly the long-lasting protective immunity. Role that has been long masked by the fact that well adapted wildtype bacteria, target these cells to prevent the clearance^18^ and facilitate reinfection.

Eosinophils are critical during immune homeostasis^19^ and lymphocyte mediated responses^28,53^. During *H. pylori* infections, bacteria manipulate eosinophils to dampen TH1 responses and promote long-term persistence infection^27^. Similarly, our results demonstrate that during infection with the *btrS*-null *B. bronchiseptica* lack of eosinophils promote long term persistence^18^ suggesting that eosinophils are required for clearance and the generation of adaptive immune responses. Increased evidence of the role of these cells during bacterial infection is mounting^54,55^, especially when looking at site specific immune responses in the mucosal gut enviroment^56^. For example, during *Pseudomonas aeruginosa* infection, eosinophilia correlates with protection against infection^57^. *Staphylococcus aureus*, activates eosinophils promoting pro-inflammatory responses associated with exacerbation of the disease^58^. In fact, tracking eosinophils numbers during disease has been proposed as a marker to evaluate antibiotherapy efficacy^59^. But why would bacteria target eosinophils to promote long-term persistent infections? Eosinophils not only have antibacterial activities themselves, but also they play important roles during the maintenance of mucosal responses^19^, due to their intrinsic relationship with lymphocites^53^,^60^,^61^. Moreover, their involvement during the formation^62,63^ and maintenance^28^ of mucosal associated lymphoid tissue is currently now elucidated. Due to their critical and important roles in maintaining the balance between anti-and auto-inflammatory responses, it makes sense that bacteria target these cells to prevent the generation of robust immune responses and protective immunity that allow not only for increased persistence but also reinfection.

Our previous published data revealed that during infection with a *B. bronchiseptica* mutant that cannot suppress host immune responses, mice present early, enhanced, and localized immune response^16^, leading to the generation of long-lasting TH17 sterilizing lung immune resposnes^17^. Based on our previous observation that eosinophils play critical roles for the clearance of infection from the lungs^18^, we investigated the mechanism by which eosinophils promote mucosal immune responses to infection with the RB50Δ*btrS* strain. We selected day 7 for most of our studies because at that day bacterial burden is the same amongst mouse strains as well as bacterial strains allowing us to identify the host factors that are driving the differences in clearance. Similar to our previously published data^16,17^ we observed an increase in the numbers of T cells in the lungs of RB50Δ*btrS* infected mice at only day 7 post-infection. Interestingly, using 2 independent mouse models we observed that the increase in T cell numbers at day 7 post challenge happened in an eosinophil dependent manner. And in the absence of eosinophils the numbers of T cells in the lungs of mice infected with the RB50Δ*btrS* strain is impaired. Eosinophils are known for their immunomodulatory roles^64^ and impact on T cell populations^65^. Moreover, eosinophils promote Th1/Th17 response upon infection with our *B. bronchiseptica* mutant. A role for eosinophils promoting TH17 responses has been previously suggested based on cytokine profiles^18^ as well as their immunomodulatory role^64,66^. It has been shown that TH17-eosinophil mediated responses are involved in the autoimmune disorders observed in patients with hyper-eosinophilic syndrome^67,68^. During *Aspergillus fumigatus* induced allergy eosinophils are also responsible for the IL-17 associated pathology observed in the lungs^69^. Moreover, during *Staphylococcus aureus* infections the number of circulating TH17 cells is increased and the authors correlate this with the fact that they found that eosinophils secrete IL17 when infected with *S. aureus* in vitro^58^. Our findings further support a role for eosinophils in promoting TH17 responses. However, these findings have been masked by the fact that wildtype *Bordetella* spp. harboring a functional *btrS* gene, manipulate eosinophils to block T cell recruitment and TH17 mucosal responses.

Even more accentuated is the increase in B cells at day 7. Eosinophils promote B cell survival, proliferation^70^, and even antibody mediated responses^61^. A role for eosinophils in activating and regulating humoral immunity has been long studied^70^. It has been shown that eosinophils promote survival and proliferation of plasma cells in the germinal centers^70^ and even in bone marrow^71,72^. Our results revealed that wildtype *B. bronchiseptica*, suppress B cell recruitment to the lungs via *btrS*-associated mechanisms and in an eosinophil dependent manner. We show that in the absence of *btrS*, there is an influx of B cells to the lungs of infected mice. However, when using two independent eosinophil deficient mouse models, the influx of B cells is significantly impaired.

Morever, our results revealed that *B. bronchiseptica* suppress/delay the formation of iBALT via btrS mediated mechanism. Our results show that the iBALT found in mice infected with the RB50Δ*btrS* strain, is formed primarily by B cells that co-localized with eosinophils where they can support B cell growth, survival, and promote humoral responses. In the absence of eosinophils iBALT was not formed. It is important to note that iBALT was more prominent in BALB/c mice. Previous research has indicated that BALB/c mice have bigger iBALT due to the increased in IL5 availiability^44^. We also observed an increase in IL5 in our unbiassed T cell secretome analysis, further supporting previously published data^44^. Our results also revealed an increased TH17 response in mice infected with RB50Δ*btrS* that was dampen in the absence of eosinophils. We investigated if eosinophils could be responsible for the increased in TH17 responses and our results revealed that eosinophils secrete IL17 in the presence of the mutant, and that in the lungs, eosinophils co-localize in the iBALT where they also promote TH17 responses by increasing the availability of IL17 within the iBALT.

Overall, these results are providing a novel role for eosinophils as promoters of pro-inflammatory responses against bacterial infections that has been masked until now due to bacterial ability to manipulate these cells. We propose a model in which *B. bronchiseptica* suppresses eosinophil effector functions via *btrS* mediated mechanism. When *btrS* is disrupted, eosinophils are recruited to the site of infection where they promote the recruitment of T and B cells to the lungs. Once there, eosinophils mediate the formation of tertiary lymphoid tissue by promoting the expression of adhesion molecules, together with cytokines and chemokines, resulting in T and B cells tertiary lymphoid tissue. Eosinophils co-localize within the aggregates where they promote TH17 mucosal responses as well as survival and proliferation of B cells and plasma cell. Altogether leading to the robust long-lasting protective immunity previously reported for this mutant. Overall, our results reveal a critical role for eosinophils during bacterial infections that has been long masked by the bacterial ability to manipulate these cells with the purpose of cause persistent infection and reinfection.

## CONCLUSION

A commonality of bacterial pathogens is that they need to overcome host immune responses. Most of the research has focused on classical phagocytes, however the role of TH2 cells during immune responses to bacterial infections is not fully understood. In this work we unravel a new role for eosinophil during the generation of TH17 protective mucosal immunity against the respiratory pathogen *Bordetella bronchiseptica*. Our results reveal that during infection with the wildtype *B. bronchiseptica*, eosinophils have no effect in the infection dynamics in the lungs because bacteria block their pro-inflammatory roles via *btrS*-immunosuppressive mechanism. However, when the bacterial sigma factor *btrS* is removed from the genetic background of *B. bronchiseptica*, suddenly eosinophils become critical for bacterial clearance from the lungs. During infection with this mutant, eosinophils promote B/T cell recruitment to the lungs, mediate TH1/TH17 responses, and importantly, promote iBALT that has a micro-TH17 environment, possibly related with the characteristic long-term protective immunity that this mutant provides against reinfection by any of the 3 classical *Bordetella* spp. including the human pathogens *B. pertussis*. Eosinophils have the ability to control immune homeostasis and TH1/TH2 balance, and our results suggest that *Bordetella* spp. manipulate eosinophils to dampen TH1/TH17 responses, suppress iBALT and overall reduce the ability of the host to clear infection and generate protective immunity.

## MATERIALS AND METHODS

### 1. Bacterial Strains and Culture Conditions

*B. bronchiseptica* strain RB50, a *btrS* knockout mutant^16^, and a *bscN* knockout^73,74^ mutant generated in a previous study were used in this study. *B. bronchiseptica* RB50 and mutants were cultured Difco Bordet-Gengou (BG) agar (BD, cat. 248200) and supplemented with 10% sheep defibrinated blood (Hemostat) with 20 μg/mL streptomycin (VWR Life sciences) or classical LB broth (Difco) as previously described^75^.

### 2. Animal Experiments

For our animal experiments we used two complementary models, one included BALB/c and the eosinophil deficient pair, ΔdblGATA-1^29^, mice originally purchased from Jackson laboratories, Bar Harbor, ME, and then breed in our facilities; and our secondary model included C57BL/6J purchased from Jackson laboratories and their eosinophil deficient pair, EPX/MBP^-/-^ mice^30^, were donated by Dr. Elizabeth Jacobsen at the Mayo Clinic. Our breeding colonies were kept under the care of the employees and veterinarians of Louisiana State University Health - Shreveport Animal Care Facility, Shreveport, LA, (AUP:20-038, AUP:22-031). All experiments were carried out in accordance with all institutional guidelines (AUP:20-0038, AUP:22-031). For our animal inoculations mice were anesthetized with 5% isoflurane and when sleep they were intranasally challenged with 30μl of PBS containing 1×10^5^ CFU/mL *B. bronchiseptica* RB50, RB50Δ*bscN* and RB50Δ*btrS*. Mice were euthanized using 5% CO_2_ followed by cervical dislocation in accordance with the humane endpoints included in our protocols.

All our animal experiments were performed in 2-3 individual experiments where we used at least 4 mice per time and condition were used in each experiment. In each figure the number of mice is indicated as n followed by the statistical test used in each individual analysis.

#### 2.1 Mouse transcriptomics

At day 7 post-inoculation, mice were euthanized according with humane endpoints and our protocol approval. Lungs were collected in 4 different beads tubes containing Trizol (Invitrogen) and kept on ice for the shortest time possible before being homogenized and frozen at −20°C. For RNA extraction, we followed the protocols recommended by the manufacturer, PureLink RNA Extraction kit, (Invitrogen) and the 4 different tubes were pooled together to have the whole lung RNA represented in one only preparation. We used 3-4 naïve animals and between 5-6 infected mice for each condition.

Twenty-four total RNA samples were assessed and processed for sequencing. Samples were quantitated with a Qubit RNA assay (ThermoFisher Scientific) and RNA quality was determined with the Agilent TapeStation RNA assay (Agilent Technologies). Libraries were prepared with the Stranded mRNA Prep, Ligation kit (Illumina). One ug of RNA was processed for each sample: mRNA was purified and fragmented. cDNA was synthesized, and 3’ ends were adenylated. Anchor sequences were ligated to each sample and limited-cycle PCR was performed to amplify and index the libraries. The average library size (330-355) was determined using an Agilent TapeStation D1000 assay (Agilent Technologies) and libraries were quantitated with qPCR (Bio-rad CFX96 Touch Real-Time PCR, NEB Library Quant Kit for Illumina). Libraries were normalized to 0.5 nM and pooled. The library pool was denatured and diluted to approximately 100 pM. A library of 2.5 pM PhiX was spiked in as an internal control. Paired-end 75 cycle sequencing was performed on an Illumina NovaSeq 6000 (Supplementary Report).

Results obtained by RNA-Seq were further validates using qRT-PCR following the recommendations of the manufacturer, Luna One-Step qRT-PCR (New England Biolabs) using the primers for each specific gene of interest (**Table 2**). qRT-PCR was performed in the CFX96 Bio-Rad. Negative controls and non-template controls were used in each qRT-PCR to exclude possible contaminations. Analysis of the results to obtain ΔΔct values was calculated after normalization with Actin and following the instructions provided in the Lune One-Step qRT-PCR kit as well as previously published protocols. All qRT-PCR were performed with at least 4 different animals, in three technical replicates each time and in our both independent animal models previously explained.

**Table 1:**
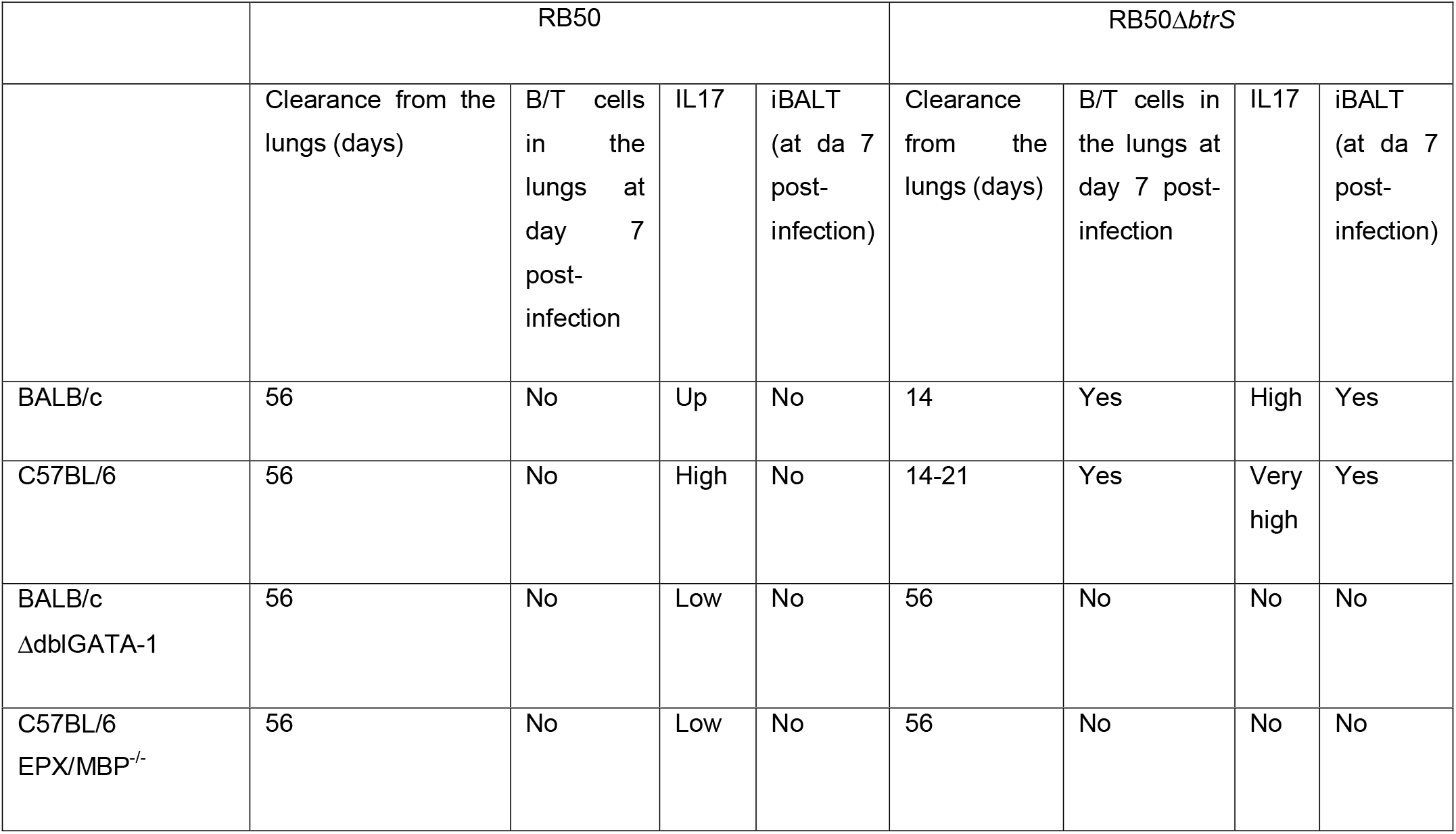
Summary of the major findings.

**Table 2:**
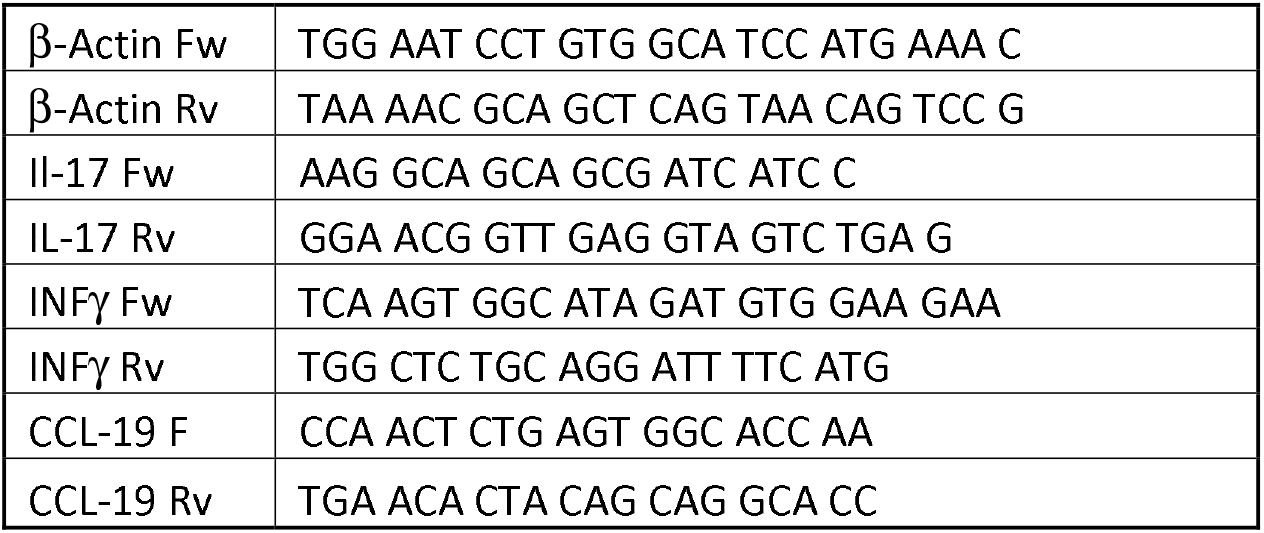
Primers used in this study

##### 2.1.1 RNA-Seq data processing

Primary analysis, including base calling and quality scoring (Supplementary Report), was performed onboard the Illumina NovaSeq 6000 (NovaSeq Control Software v1.7.5; RTA v3): Samples were de-multiplexed, the anchor sequences were removed (the first 9 cycles of sequencing were trimmed), and FASTQ files were generated. FASTQ files were downloaded from BaseSpace Sequencing Hub and consolidated. Reads were aligned to the mouse genome (GRCh38.p28, GenBank Assembly GCA_000001635.9) using STAR__2.5.31 and counted using RSEM1.3.0.

###### Galician Supercomputational Centre

FASTQ files were processed using the computational resources of the Galician Supercomputational Centre (CESGA). Adaptors and low-quality bases were removed using Trimmomatic (version 0.38). Index was generated using Rsem software (version 1.3.1) utilizing Mus musculus reference genome version GRCm39and gene transfer format (.gtf) annotation version GRCm39.105 from Ensembl. Reads were aligned using STAR (version 2.7) and expression values were calculated using Rsem (version 1.3.1).

Differential gene expression was evaluated using DESeq2 R package (version 1.34.0). Benjamini-Hochberg correction was used to obtain adjusted p-values. Shrunken log2 fold changes were obtained using ‘ashr’ shrunkage^76^. Genes with adjusted p-value < 0.05 and absolute log2 fold change ≥ 1.5 were considered significant in terms of differential expression.

Gene Set Enrichment Analysis (GSEA) and over-representation (OR) test for differentially expressed genes were performed using Clusterprofiler (version 4.2.2)^77,78^ and Gene Ontology – Biological Process collection for mouse. Benjamini-Hochberg correction was used to obtain adjusted p-values. Redundant terms in over-representation analysis were reduced using the simplify function, cutoff=0.6.

#### 2.2 Flow cytometry analysis

Flow data was obtained in 2 individual biological replicates containing 4-6 mice per time and condition each time. Following inoculation, mice were euthanized at day 7 post-infection following our approved protocol recommendations. Lungs were collected in 5 ml of cold PBS and were kept on ice. To obtain single cell suspensions we followed the guidelines and protocol of GentleMacs Dissociator, (Milteny). Following single cell isolation, we procedure with our standard processing and staining protocol (Table XX), including single beads staining and fluorescence minus one in each staining. Samples were acquired in our NovoCyte flow cytometer (Agilent), and gating strategy is shown in supplementary material. All immunology experiments were performed at the Immunophenotyping core, LSU Health Shreveport.

#### 2.3 T cell secretome analysis

Following inoculation with RB50 or RB50Δ*btrS*, groups of 4 mice were euthanized. Lungs were collected in 5 ml of cold PBS and were kept on ice. To obtain single cell suspensions we followed the guidelines and protocol of GentleMacs Dissociator, (Milteny). We used the single cell preparation to isolate T cells using negative selection magnetic beads, MojoSort™ Mouse CD3 T Cell Isolation Kit (BioLegend). T cells were then seed in a 96 well plate containing 10^6^ cells per well in a volume of 200 μl with RPMI (Gibco) containing 10%FBS (Gibco) and 50μM β-mercaptoethanol (Sigma). We then added 25 μl of beads CD3/CD28 (Invitrogen) and incubated at 37°C and 5% CO2 for 48 hours. Following incubation, we loaded the bulk cartridge CodePlex Secretome (IsoPlex). All immunology experiments were performed at the Immunophenotyping core, LSU Health Shreveport.

#### 2.4 Pathology analysis

Pathology experiments were performed in 2-4 biological replicates containing at least 4 mice per time and condition each time. The total number of mice was at least 6 for all the experiments done in BALB/c and ΔdblGATA-1; and at least 4 for all the experiments performed with C57BL/6J and EPX/MBP^-/-^ mice. For all pathology experiments, mice were euthanized and perfused intratracheally with sterile PBS and 4% paraformaldehyde (PFA). Lungs were subsequently, fixed overnight in 10 mL of 4% PFA prior to processing, and paraffin embedding. Lungs are processed, paraffin-embedded, sectioned in 0.5 μm thick slices, and placed on glass slides by the COBRE Histology Core at LSU Health Shreveport. All our microscopy analysis were performed in three sections per mouse; and we collected at least 10 areas per section.

##### 2.4.1 Immunofluorescence and Image Acquisition/Analysis

To perform our immune staining, we followed protocols previously published by our collaborators^79^. Briefly, slides with PFA fixed and paraffin-embedded tissues were deparaffinized in consecutive washes in xylene, followed by rehydration, washes, and citrate buffer solution for antigen retrieval^79,80^. Antigen retrieval wash was done by steaming slides in a citrate buffer with a pre-heated Instant Pot pressure cooker for 10 minutes at medium heat. After cooling down and washes, PBS with 0.2% Triton-X 100 (PBST) was used to permeabilize the tissues. After bocking for 90 minutes in 13 mL of 0.15% Bovine Serum Albumin (BSA) with Normal Goat Serum and PBST, slides underwent PBST washes prior to staining. Staining was performed using the appropriate primary antibodies diluted with 0.15% BSA in PBST (**Table 3)** and incubated overnight. The next day, secondary antibodies were used prior to Hoechst 33342 staining. Slides coverslips were mounted to the sections with ProLong™ Glass Antifade Mountant and dried in the dark at room temperature until later analysis. Immunofluorescent images of the lungs were captured using a Keyence BZX-800 microscope. Image analysis was conducted using the Keyence Image Analysis Software. Briefly, for analysis of each immunofluorescence assay a fluorescence intensity threshold was set and used to detect Hoechst stain lung cells that were also positive for each target antibody. Each group of images analyzed in this manner was done in batch analysis using the same threshold settings.

**Table 3:** Antibodies and immunofluorescence reagents used in this study

| Primary Antibodies | Vendor | Catalog # |
| --- | --- | --- |
| Alexa Fluor® 488 anti-mouse CD3 Antibody | Biolegend | 100210 |
| CD45R (B220) Monoclonal Antibody (RA3-6B2), eBioscience™ | Invitrogen | 14-0452-82 |
| Bone Marrow Proteoglycan (PRG2) Antibody or Major Basic Protein (MBP) | Abbexa | Abx101775 |
| Rabbit anti-mouse LYVE1 antibody | Abcam | Ab149117 |
| Secondary Antibodies | Vendor | Catalog # |
| Alexa Fluor® 594 donkey anti-rat IgG (H+L) | ThermoFisher Scientific | A21209 |
| Alexa Fluor® 488 goat anti-rabbit IgG (H+L) | ThermoFisher Scientific | A11008 |
| Donkey Anti-Rabbit IgG Antibody, HRP conjugate | Millipore-Sigma | AP182P |
| Goat Anti-Rat IgG Antibody, HRP conjugate | Millipore-Sigma | AP136P |
| Other IF Reagents | Vendor | Catalog # |
| Hoechst 33342 Solution | BD Biosciences | 561908 |
| Opal 520 Reagent Pack | Akoya Biosciences | FP1487001KT |
| Opal 570 Reagent Pack | Akoya Biosciences | FP1488001KT |
| Opal 650 Reagent Pack | Akoya Biosciences | FP1496001KT |

### 3. Eosinophils differentiation and infection

BALB/c mice were euthanized accordingly with our protocol guidance. We cleaned the femur and tibia. We flashed the inside of the bones with RPMI to obtain Bone marrow progenitors. To differentiate them into eosinophils we followed previously published protocols^81^. Briefly, Single cell suspensions of bone marrow progenitors were incubated in eosinophils base media containing 100ng/ml rmFLT3L (PrepoTech) and 100 ng/ml rmSCF (PrepoTech) for 4 days. After that the cells were maintained in eosinophil base media supplemented with 10 ng/ml IL5 (PrepoTech) up to 16 days. Percentage of differentiation was evaluated by flow cytometry and/or cytochemical staining^81^.

For the infection, multiplicity of infection (MOI) was calculated in function of the number of eosinophils per well, confluency was not considered, as eosinophils do not attach. MOI of 0.1, 1, and 10 were used on our assays as indicated in each figure. Bacteria were resuspended in eosinophils base media to perform the infections. After adding bacteria, we spin down the plates 300 g for 5 minutes and we started time cero. All the assays were done in at least 3 independent biological replicates containing 6 technical replicates each time.

### 4. LegendPlex

Supernatant of the in vitro infections was collected at 4 hours post-infection and kept at −20 until its used. For the assay we followed the recommendations for LegendPlex of the manufacturer (Biolegend). We used at least 4 independent biological replicates all in technical duplicates as indicated in each figure legend. For the analysis we used the cloud software recommended legenplex.qognit.

### 5. Statistical analysis

All experiments were performed in three independent biological replicates. The exact number of mice and technical replicates is indicated in each figure legend. For animal experiments we performed Two-Way ANOVA analysis, while qRT-PCR experiments were analyzed using One-Way ANOVA analysis. Similarly, all the data that was normally distributed was analyzed using One-way and Two-way ANOVA test as indicated in each figure legend. When ANOVA was not an option we performed t-Student Tests. All graphs and data were analyzed using the GraphPad V9.0. A p value ≤0.05 was considered statistically significant. In the figures the asterisks correspond with * p≤0.05, ** p≤0.01, *** p≤0.001, and **** p≤0.0001.

## ACKNOWLEDGEMET

We would like to acknowledge the support of the funding bodies: NIH COBRE award 1P20-GM134974-0181. Louisiana Board of Regents LEQSF(2022-25)-RD-A-33. Center of Excellence for Arthritis and Rheumatology intramural award. Intramural Research Council Seed package support from LSU Health, Shreveport. Start-up package from LSU Health, Shreveport.

We would like to acknowledge the two centers funded by our COBRE P20-GM134974 The immunophenotyping has helped with the antibody titration, staining, data analysis and all the immunology. The modelling core has helped with data analysis and the generation of figures. We would also like to acknowledge Martin Sapp for his ideas, brainstorming, and support during the editing process.

## Author contribution

NJF performed in vitro experiments, analyzed data, contributed to writing and editing. AM brainstorming, analyzed data, contributed to the writing and editing. AGF performed brainstorming, analyzed data, contributed to writing and editing. SB performed experiments, analyzed data, and contributed to editing. CR performing experiments and analyze data. Emily Cox performing experiments and analyze data. JW analyze data. RS analyze data, brainstorming, and contributed editing. MW brainstorming and contributing to writing and editing. MCG conceptualized the project, designed experiments, performed experiments, analyzed data, writing and editing, obtained funding, supervised.

