## supplementary figures for "*Bordetella* spp. block eosinophil recruitment to suppress lung iBALT formation"

^2^CINBIO, Universidade de Vigo, Immunology group, 36310 Vigo, Spain. Instituto de Investigación Sanitaria Galicia Sur (IIS Galicia Sur)

^3^Immunophenotyping Core. Center for Applied Immunology and Pathological Processes. Department of Microbiology and Immunology, LSU Health Science Center, Shreveport. 71103, LA, USA

^4^Modelling Core. Center for Applied Immunology and Pathological Processes. Department of Microbiology and Immunology, LSU Health Science Center, Shreveport. 71103, LA, USA

Corresponding: Monica Cartelle Gestal, PhD. //

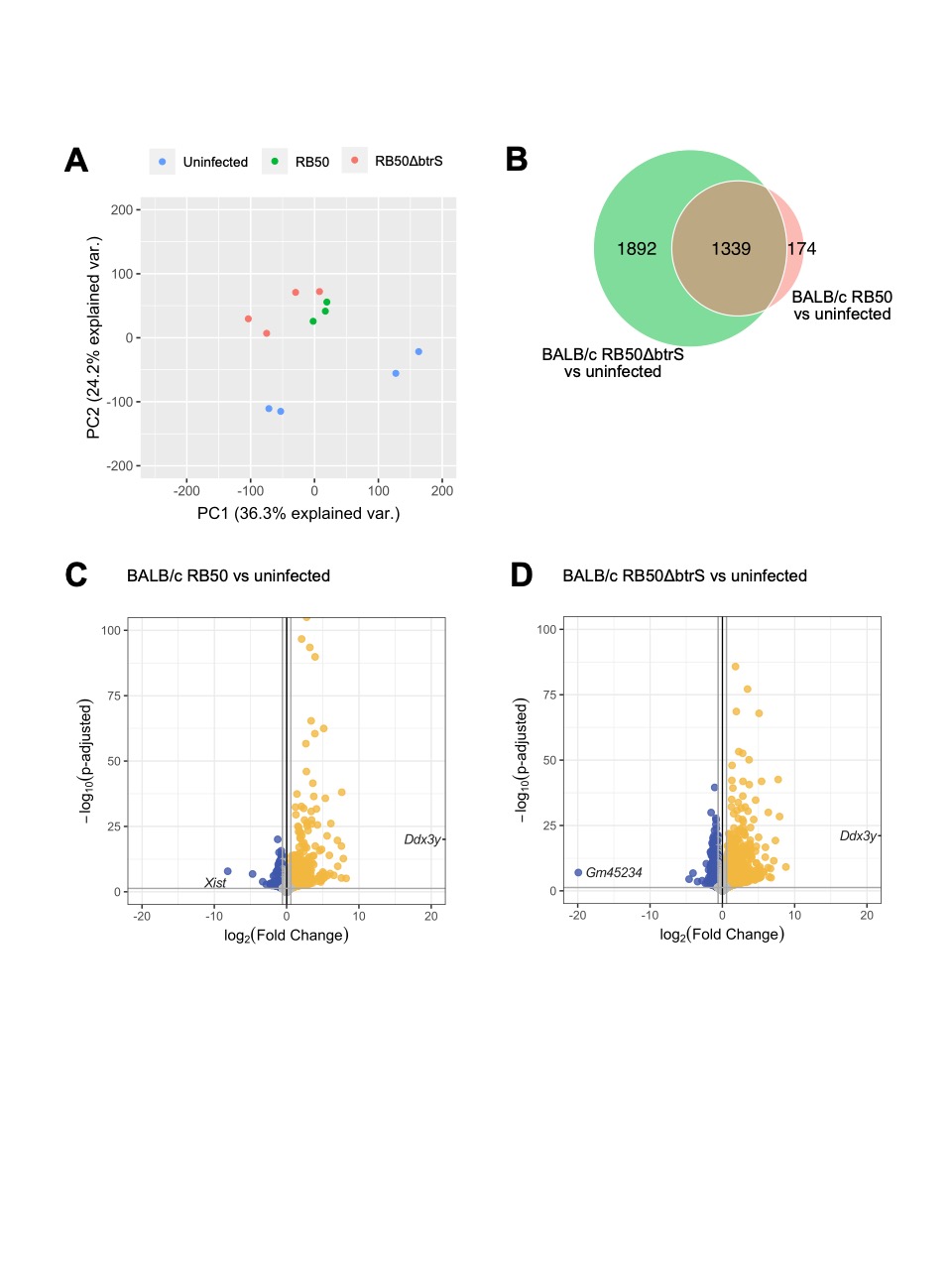

**Figure S1: Infection with RB50Δ*btrS* promote more transcriptional changes in the lungs of BALB/c mice.** PCA of normalized RNA-seq read counts in Transcripts Per Million (TPM) of BALB/c mice. Sample to sample distances along the PC1 and PC2 were depicted for all samples. Samples were labelled depending on the mouse strain and colored depending on type of infection. (B) Venn-Diagram representation of the number of differentially expressed genes compared to uninfected controls. (C,D) Volcano plot representation of BALB/c infected with RB50 or RB50Δ*btrS* versus uninfected BALB/c. Gray dots represent all genes, whereas colored dots depict genes differentially expressed genes (blue: downregulated, yellow: upregulated). Names of the top upregulated and downregulated genes ordered by fold change are plotted (n=3-5 mice per group).

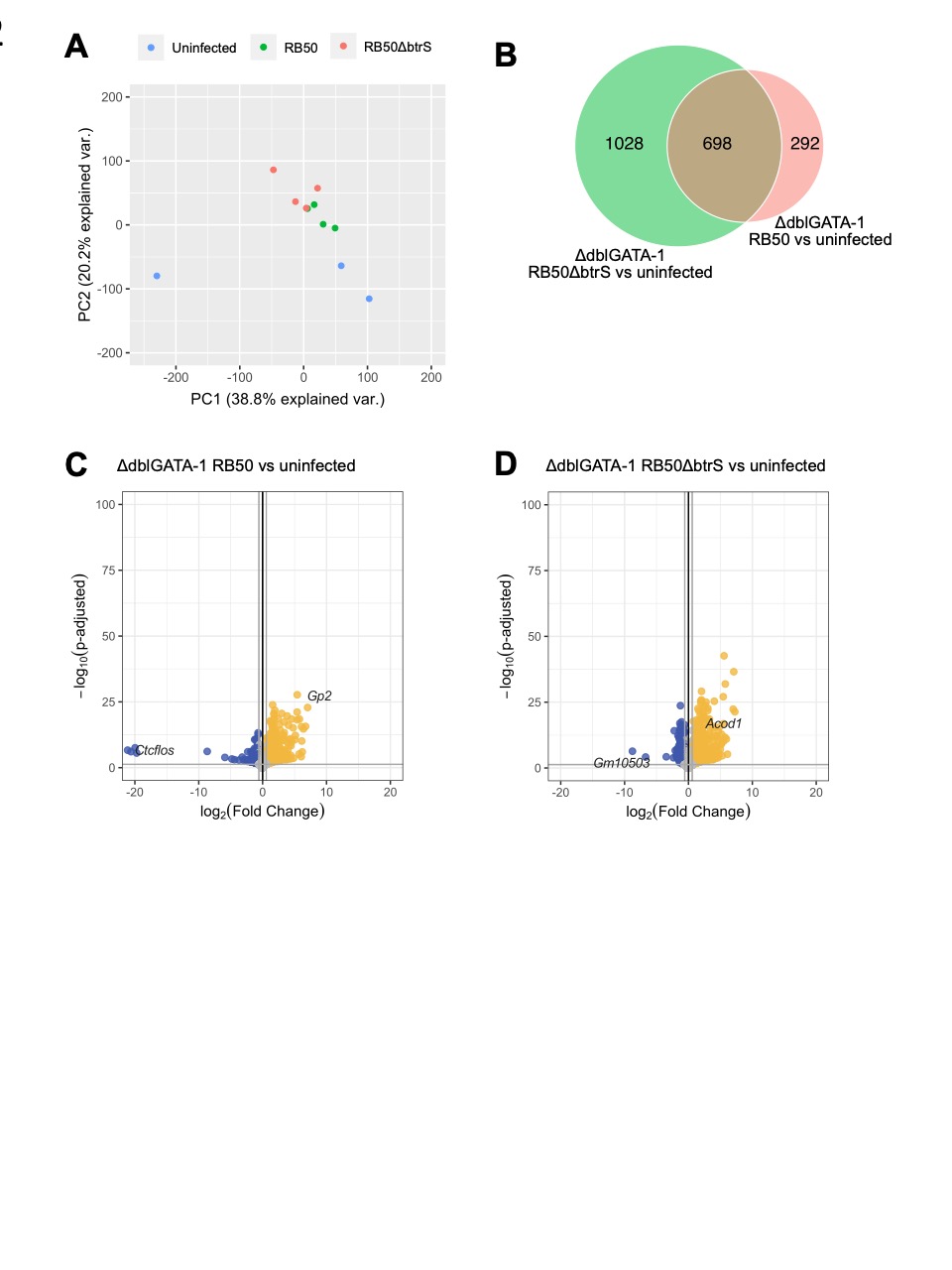

**Figure S2: Infection with RB50Δ*btrS* promote more transcriptional changes in the lungs of ΔdblGATA-1 mice.** PCA of normalized RNA-seq read counts in Transcripts Per Million (TPM) of dblGATA-1 mice. Sample to sample distances along the PC1 and PC2 were depicted for all samples. Samples were labelled depending on the mouse strain and colored depending on type of infection. (B) Venn-Diagram representation of the number of differentially expressed genes compared to uninfected controls. (C,D) Volcano plot representation of ΔdblGATA-1 infected with RB50 or RB50Δ*btrS* versus uninfected ΔdblGATA-1 Gray dots represent all genes, whereas colored dots depict genes differentially expressed genes (blue: downregulated, yellow: upregulated). Names of the top upregulated and downregulated genes ordered by fold change are plotted (n=3-5 mice per group).

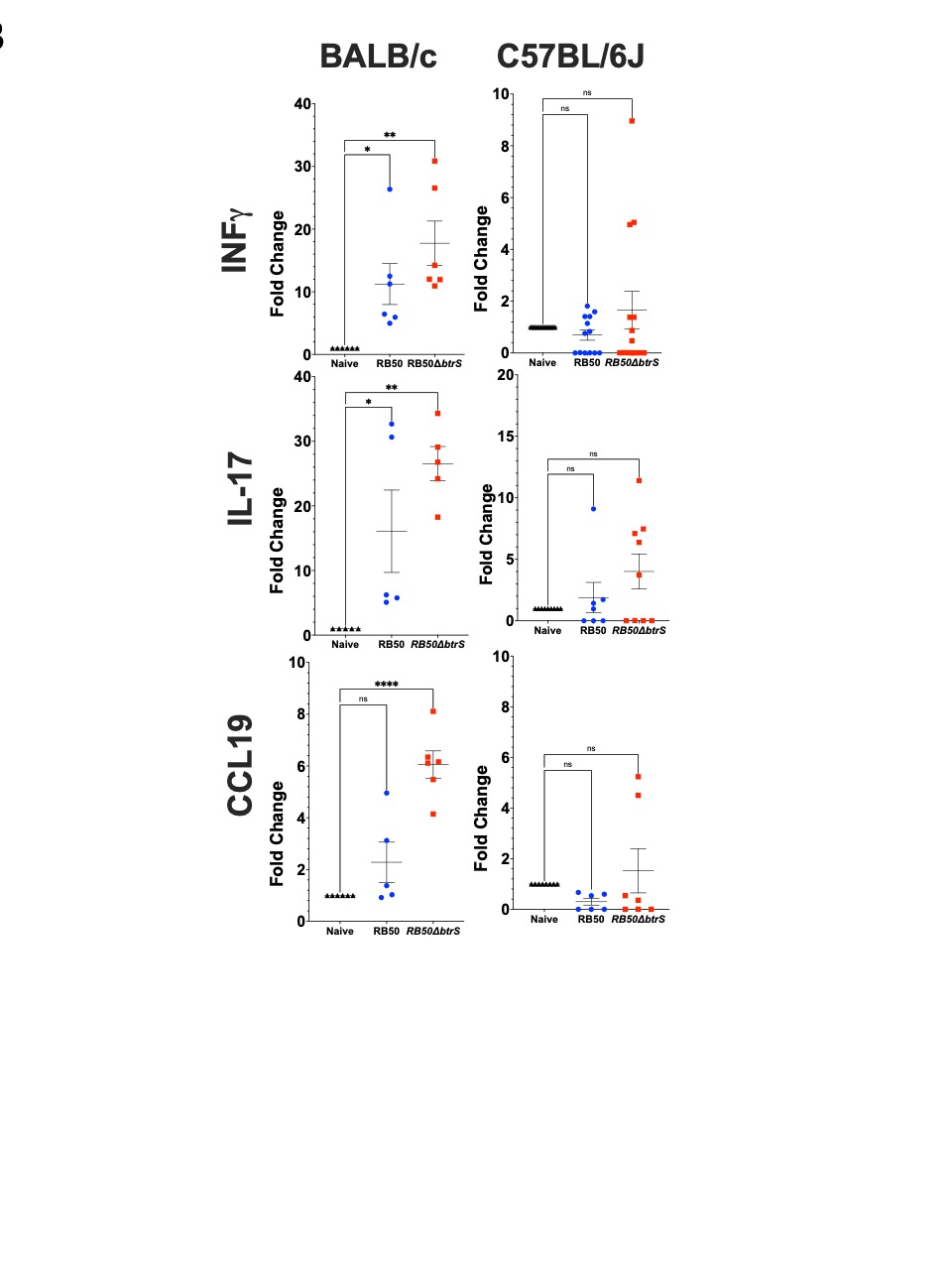

**Figure S3: Infection with RB50Δ*btrS* promote pro-inflammatory responses in BALB/c and C57BL/6J mice.** RNA from the lungs was utilized to perform qRT-PCR and measure mRNA levels using actin as a normalizer of our data. Levels of mRNA for INFγ, IL-17, and CCL19, were measured in uninfected (black), infected with RB50 (blue) and infected with RB50Δ*btrS* (red) mice at day 7 post-infection. One-way ANOVA was used to determine statistical significance. * p≤0.05, ** p≤0.01, and **** p≤0.0001 (n= 4-8 mice per group).

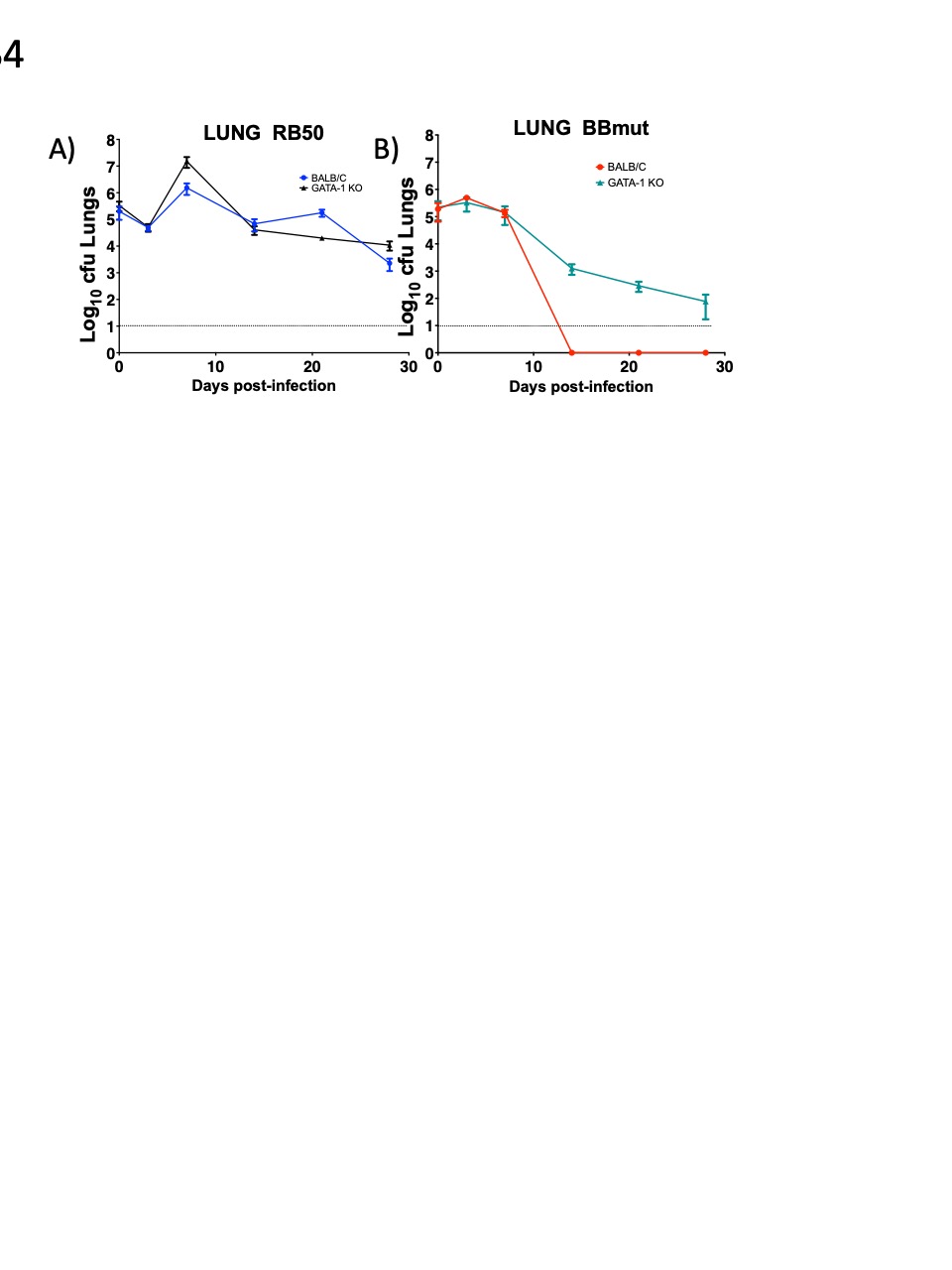

**Figure S4: Eosinophil deficient mice fail to clear infection by RB50Δ*btrS* but not by RB50.** Groups of mice were infected with RB50 or RB50Δ*btrS* and at different post-infection times, lungs bacterial burden was enumerated by plating serial dilutions. (A) Infection with RB50 in BALB/c mice (blue) results in the same dynamic of infection in ΔdblGATA-1 mice (black). (B) Infection with RB50Δ*btrS* in BALB/c mice (results) results more rapid clearance than in ΔdblGATA-1 mice (green) (n=4-12 mice per group).

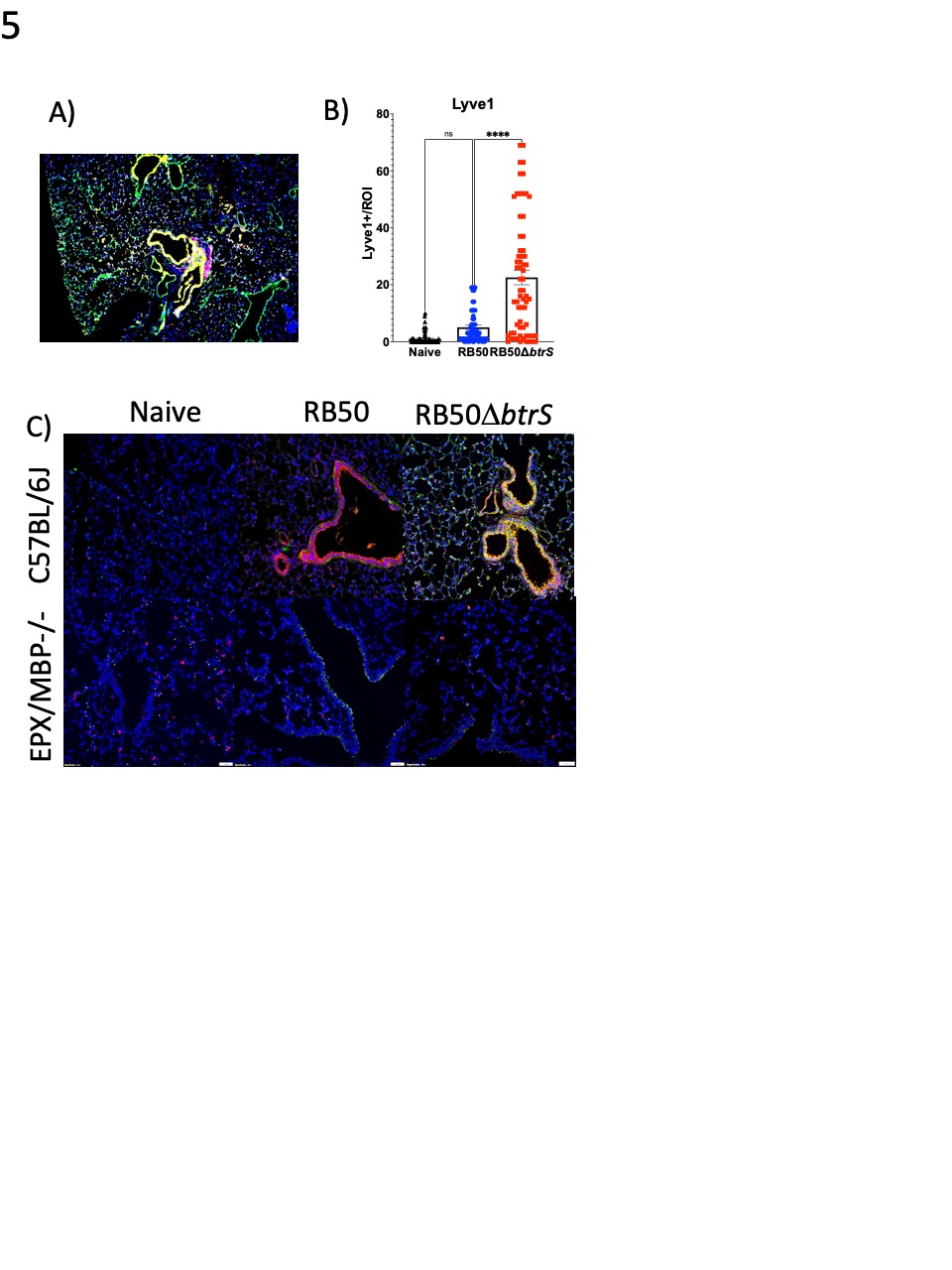

**Figure S5: Infection with RB50Δ*btrS* promotes the formation of iBALT.** Groups of mice were infected with RB50 or RB50Δ*btrS* and at day 7post-infection, lungs were collected for immunohistochemistry. (A) Lungs of RB50DbtrS infected mice were stained for Hoechst (blue), B220 B cells (red) and Lyve-1 endothelial marker (green). (B) Lyve-1 positive signal was measured in three sections per mouse and 10 regions of interest (n= 4 mice per group). Groups were uninfected (black), infected with RB50 (blue), and infected with RB50Δ*btrS* (red). (C) C57BL/6J and EPX/MBP^-/-^ mice were intranasally challenged with PBS (naïve), RB50, and RB50Δ*btrS*. At day 7 post-infection lungs were stained with Hoechst (blue), B220 B cells (red) and CD3 T cells (green). Three sections per mouse and 10 regions of interest per section, were used to evaluate iBALT (n= 6 mice per group).
